# Disrupted remapping of place cells and grid cells in knock-in model of Alzheimer’s disease

**DOI:** 10.1101/815993

**Authors:** Heechul Jun, Shogo Soma, Takashi Saito, Takaomi C Saido, Kei M Igarashi

**Affiliations:** Department of Anatomy and Neurobiology, University of California, Irvine; Center for the Neurobiology of Learning and Memory, University of California, Irvine; Institute for Memory Impairments and Neurological Disorders, University of California, Irvine; Lab for Proteolytic Neuroscience, RIKEN Center for Brain Science, Japan; Japan Science and Technology Agency

## Abstract

Patients with Alzheimer’s disease (AD) frequently suffer from spatial memory impairment and wandering behavior, but brain circuits causing such symptoms remain largely unclear. In healthy brains, spatially-tuned hippocampal place cells and entorhinal grid cells represent distinct spike patterns in different environments, a circuit function called “remapping” that underlies pattern separation of spatial memory. We investigated whether knock-in expression of mutated amyloid precursor protein deteriorates the remapping of place cells and grid cells. We found that the remapping of CA1 place cells was disrupted although their spatial tuning was only mildly diminished. Grid cells in the medial entorhinal cortex (MEC) were impaired, sending severely disrupted remapping signals to the hippocampus. Furthermore, fast gamma oscillations were disrupted in both CA1 and MEC, resulting in impaired fast gamma coupling in the MEC→CA1 circuit. These results point to the link between grid cell impairment and remapping disruption as a circuit mechanism causing spatial memory impairment in AD.

## Main Text

Spatial memory impairment such as wandering behavior is one of the most troublesome symptoms in Alzheimer’s disease (AD), occurring in more than 60% of AD patients (Hope et al., 1994). Despite recent molecular and cellular findings in AD research, it is still largely unclear how deterioration of brain circuit function causes spatial memory loss in AD. In healthy brains of both humans and rodents, grid cells in the medial entorhinal cortex (MEC) and place cells in the hippocampus form a neural circuit that is critical for spatial memory and navigation (Fig. S1) (Scoville and Milner, 1957; O’Keefe and Dostrovsky, 1971; Morris et al., 1982; Ekstrom et al., 2003; Fyhn et al., 2004; Steffenach et al., 2005; Doeller et al., 2010). In this circuit, spatially-tuned grid cells in the superficial layer of MEC send spatial signals to the hippocampus to form spatially-tuned place cells (Moser et al., 2015). These grid cells and place cells have an ability to store multiple combinations of firing patterns when subjects enter into different environments, a function called “remapping” (Muller and Kubie, 1987; Fyhn et al., 2007; Alme et al., 2014; Kyle et al., 2015). Because remapping of place cells and grid cells can provide subjects with distinct combinatorial codes for multiple environments, it provides support for the circuit mechanism for the pattern separation of spatial memory (Colgin et al., 2008; Yassa and Stark, 2011). However, no previous studies have investigated remapping of place cells and grid cells in AD subjects.

In this study, we specifically asked two questions: (1) Is remapping of hippocampal place cells deteriorated in AD subjects? (2) If so, is place cell deterioration linked to impairment of entorhinal grid cells? Alternatively, does place cell deterioration occur autonomously inside the hippocampus and independent from grid cells? To this end, we investigated the remapping of hippocampal place cells and entorhinal grid cells in a novel amyloid precursor protein knock-in (APP-KI) mouse we recently developed (Saito et al., 2014). Previous transgenic AD models express an excessive amount of mutated genes from multiple copies of transgenes in both the entorhinal cortex and the hippocampus, making it difficult to compare pathophysiology between the two brain regions. By contrast, in our knock-in model, mutated APP protein is expressed from only two copies of mutated human APP gene, as is the case for human AD patients. Therefore, relative pathological progressions between the hippocampus and the entorhinal cortex of APP-KI mice are expected to recapitulate those from AD patients, allowing us to compare the pathophysiology between the hippocampus and the entorhinal cortex. Because APP-KI mice show severe spatial memory impairments (Masuda et al., 2016), we investigated the remapping of place cells and grid cells underlying the spatial memory impairments.

## Results

### Hippocampal CA1 neurons showed severely disrupted remapping, but mildly diminished spatial tuning in APP-KI mice

In AD patients, pathophysiology starts with amyloid-β (Aβ) plaque deposition from the expression of APP, which slowly leads to Tau pathology and deterioration of brain circuit functions in the course of more than 20 years (Sasaguri et al., 2017). The circuit deterioration ultimately leads to memory impairment, with which patients are diagnosed with AD. To replicate the slow and cumulative effect of pathological APP on brain circuit functions in mice, we recently generated a knock-in mouse model by manipulating the endogenous murine APP gene into the mutated human APP^NL-F-G^ gene (Saito et al., 2014). This knock-in manipulation causes Aβ deposition starting at 4-mo and severe spatial memory impairment at 6-mo (Saito et al., 2014; Masuda et al., 2016). To investigate place cells after memory impairment, we implanted a recording device containing 64 channels of electrodes in APP-KI mice and recorded place cells after 7-mo (7-13 mo, Fig. 1A-B **and** S2-S3; no change observed in place cells and grid cells from APP-KI mice across this age period, Fig. S4). Because the APP-KI mouse is on the C57BL/6 background, we used 7-13 mo C57BL/6 mice as a control (referred to as WT mice). No behavioral difference was observed between APP-KI and WT mice (Fig. S5). To avoid biased sampling toward spatially tuned cells that can be fewer in APP-KI mice, we collected all recorded principal neurons in our data pool (n=105 neurons from n = 10 APP-KI mice; n=113 neurons from n = 10 WT mice). No difference was observed in the distribution of putative principal neurons and interneurons between WT and APP-KI mice (Fig. S6). Neural activities of CA1 neurons were first recorded in a 1×1 m open field to evaluate their spatial tuning and place cell properties (Fig. 1C). Although our WT mice were between the age of 7-13 months, a good amount of CA1 neurons in WT mice showed spatially tuned spike activities (Fig. 1D). CA1 neurons in APP-KI mice also showed spatially tuned spike activity, but exhibited less pronounced firing peaks (Fig. 1E). The degree of spatial tuning for individual CA1 neurons was quantitatively assessed using the *spatial information* score (Skaggs et al., 1993) (Fig. 1F-H). The analysis revealed that APP-KI mice have neurons with less spatial tuning (p<0.001, Kolmogorov-Smirnov (KS) test), with lower mean spatial information than those from the CA1 neurons recorded in the WT mice (p<0.001, Wilcoxon rank sum test; detailed statistic results are shown in Supplementary Tables S2-S14). However, mean firing rates, field size and firing field number of CA1 neurons did not differ between WT and APP-KI mice, suggesting the spatial tuning of CA1 neurons in APP-KI mice was only mildly deteriorated (Fig. S7). We then defined place cells using the spatial information score (Langston et al., 2010). Neurons were classified as place cells if their spatial information score was larger than 0.63, which was the 95th percentile of a distribution of spatial information score based on shuffled data from WT mice (Fig. 1F-G **and** S8). This revealed that proportion of place cells decreased down from 50% in WT mice to 31% in APP-KI mice (p<0.001, binomial test; Fig 1G-H). Among these place cells, we did not observe any differences in their properties recorded from the open field task between APP-KI and WT mice (Fig. S8).

**Figure 1.**
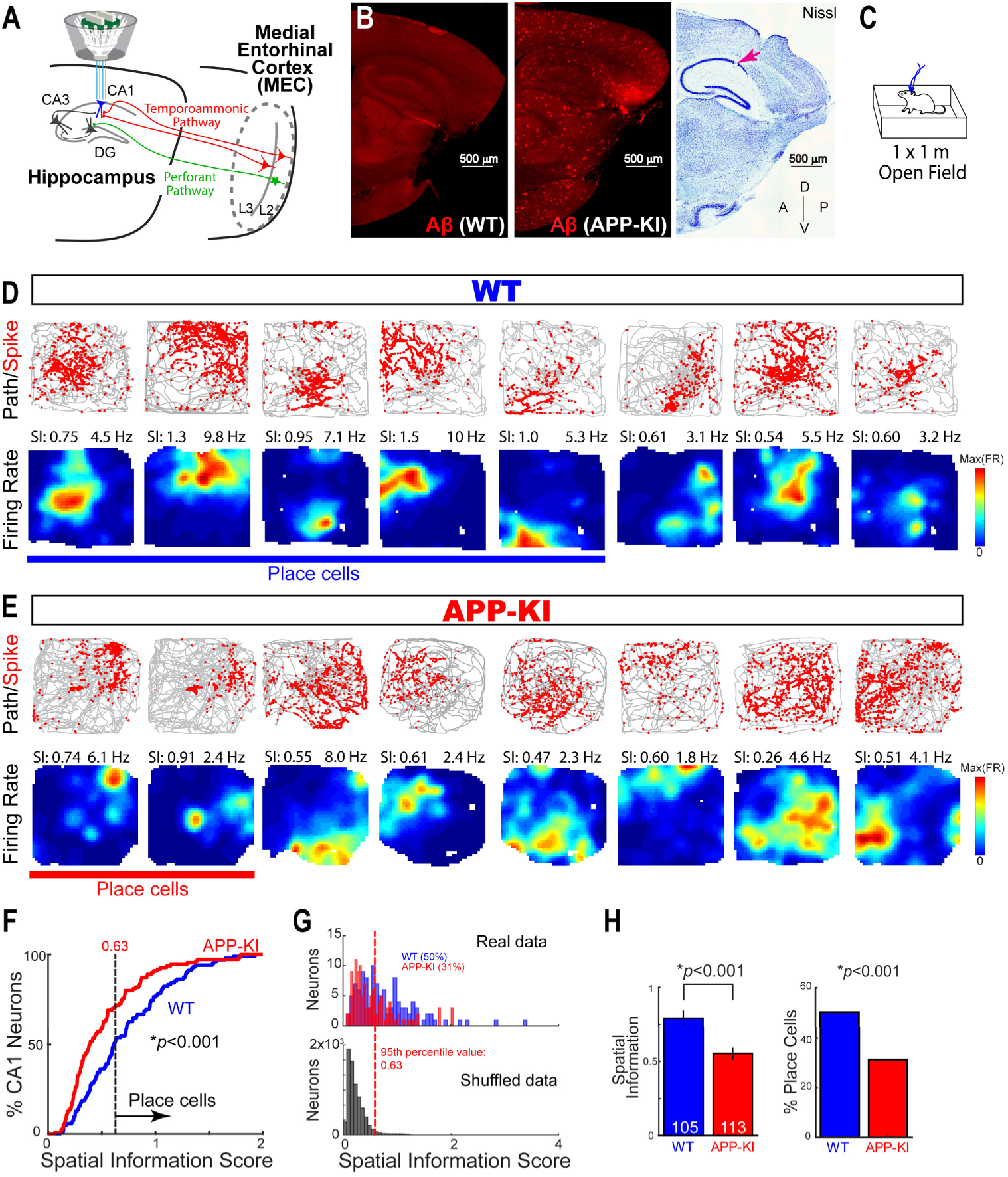
Hippocampal CA1 neurons, including place cells, showed mildly diminished spatial tuning in APP-KI mice. (**A**) A 64-channel recording device was targeted to the CA1 region of the hippocampus. (**B**) Sagittal sections with anti-Aβ immunostaining from 12-mo WT mouse (left) and 12-mo APP-KI mouse (middle), and with Nissl staining showing a representative recording position in CA1 (right, red arrow). A, anterior; P, posterior; D, dorsal; V, ventral. Scale bars, 500 µm. (**C**) 1×1 m open field for testing spatial tuning of CA1 neurons. (**D**) Eight representative CA1 neurons in WT mice. Top: Red dots and gray lines denote spike position and animal trace in the 1×1 m open field, respectively. Bottom: Firing rate map. Color is scaled with maximum firing rate (Hz) shown at top right of each rate map. Spatial information (SI) score is shown at top left. (**E**) Eight representative CA1 neurons as in (D), but from APP-KI mice. (**F**) Cumulative distribution plot for spatial information score of CA1 neurons in WT mice (blue, n=105) and APP-KI mice (red, n=113). A threshold of 0.63, obtained from 95^th^ percentile of shuffled WT data, was used to define place cells. p<0.001, KS test. (**G**) Top, distribution of spatial information scores calculated from firing rate distributions of CA1 neurons in the open field task (real data). Bottom, distribution of shuffled data based on 100 permutations per cell. Red dashed line indicates 95th percentile value (chance level) for a distribution based on all permutations. Percentage of cells that exceeded chance level is shown on top. 50% of CA1 neurons in WT mice and 31% of CA1 neurons in APP-KI mice passed this chance level, and they were considered as place cells. (**H**) Left: Mean spatial information score of CA1 neurons in WT mice (n=105) and APP-KI mice (n=113). p<0.001, Wilcoxon rank sum test. Right: Percentage of neurons defined as place cells. p<0.001, binomial test.

To investigate the remapping of CA1 neurons in APP-KI mice, we then trained animals to run back and forth in two 1m linear tracks (Tracks A and B) with distinct colors and textures (Fig. 2A, see experimental protocol). In a daily session, animals made 10 laps for each track in the following series: Track A, Track B, Track B and then back again in Track A. On the fifth day of the training, CA1 neurons in WT mice showed remapping across Track A and Track B, with distinct firing patterns between the two tracks (Fig. 2B, also known as “global remapping” (Leutgeb et al., 2005b)). However, with the same training condition, CA1 neurons in APP-KI mice showed unchanged firing patterns between Track A and Track B (Fig. 2C). Population vector (PV) correlation analysis (Leutgeb et al., 2005b) was used to assess the degree of remapping between Track A and Track B (Fig. 2D-E). A PV correlation of −1 indicates that the recorded population of CA1 neurons showed completely distinct firing patterns (that is, perfect remapping), whereas a PV correlation of 1 indicates exactly identical firing patterns (that is, no remapping at all). PV correlation analysis revealed that WT mice showed good remapping with low PV correlation between Track A and Track B (Fig. 2E). However, APP-KI mice exhibited PVs with significantly higher correlation, indicating that the remapping of CA1 neurons was significantly disrupted in APP-KI mice (p<0.001, KS test, Fig. 2E). Mean PV correlation in APP-KI mice was significantly higher than that of WT mice (p<0.001, Wilcoxon rank sum test; Fig. 2F). Disrupted remapping was observed independently of the direction of animals’ runs in the tracks (Fig. S10). Remapping of the neurons defined as place cells was also significantly disrupted (Fig. S8). We also assessed the degree of remapping in individual neurons using the *spatial correlation* score between Track A and Track B (Leutgeb et al., 2005b), confirming that CA1 neurons showed disrupted remapping in APP-KI mice compared to that from WT mice (Fig. 2G **and** S11). By contrast, activity patterns of place cells between the two recordings in Track A were similar, with no difference in PV correlations between WT and APP-KI mice, indicating that stability of spatial tuning in CA1 neurons was spared in APP-KI mice (p>0.05, KS test, Fig. 2E; p>0.05, Wilcoxon rank sum test, Fig. 2F). Together, these results demonstrate that CA1 principal neurons from APP-KI mice exhibit disrupted remapping for different spatial environments, while the spatial tuning of CA1 neurons is only mildly deteriorated.

**Figure 2.**
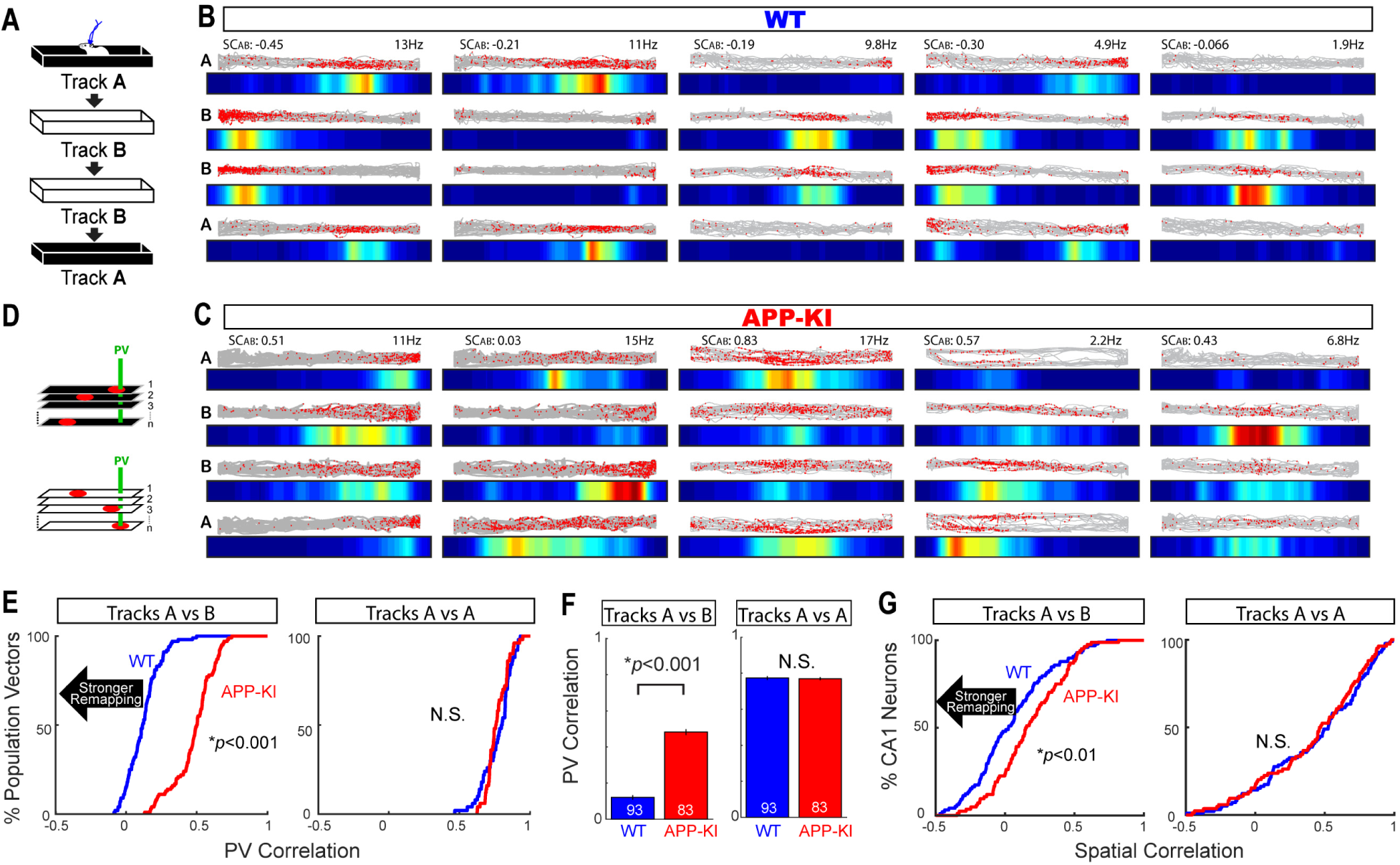
Hippocampal CA1 neurons in APP-KI mice showed disrupted remapping. (**A**) Remapping was tested in 1-m linear Track A and Track B with distinct colors and textures. (**B**) Five representative CA1 neurons in WT mice recorded in Tracks A, B, B, and A (from Top to Bottom). Red dots and gray lines at the top of each track denote spike position and animal trace in the 1 m linear tracks, respectively. Bottom color map denotes firing rate map. Color is scaled with maximum firing rate (Hz) shown at top right. Spatial correlation between Track A and Track B (SCAB) is shown at top left. (**C**) Five representative CA1 neurons from APP-KI mice are shown as in (B). (**D**) For evaluating the degree of remapping, a population vector (PV) was constructed for each spatial bin (green) in Track A and Track B, and their correlation was analyzed (PV correlation). (**E**) Cumulative distribution plots for PV correlation between Track A and Track B (left) and between two recordings in Track A (right). Lower PV correlation denotes stronger remapping (black arrow). p<0.001, KS test. (**F**) Left: Mean PV correlation between Track A and Track B. APP-KI mice showed significant increase (that is, weaker remapping) compared to that of WT mice. p<0.001, Wilcoxon rank sum test. Right: Mean PV correlation between two recordings in Track A did not differ. p>0.05, Wilcoxon rank sum test. (**G**) Left: Cumulative distribution plots of spatial correlation between Track A and Track B from CA1 neurons during remapping task. Significant increase in APP-KI mice compared to that of WT mice indicates disrupted remapping. p<0.01, KS test. Right: Cumulative distribution plot of spatial correlation between two recordings in Track A. p >0.05, KS test.

### Diminished fast gamma oscillations, but not slow gamma oscillations in the hippocampal CA1 of APP-KI mice

CA1 neurons in the hippocampus receive spatial inputs from hippocampal CA3 via the Schaeffer collateral as well as from the MEC via temporoammonic pathway (Witter and Amaral, 2004). We next asked whether these inputs contribute to disrupted remapping in CA1. We leveraged findings from previous healthy animal studies that identified two distinct types of gamma oscillations (Bragin et al., 1995; Colgin et al., 2009) (Fig. 3A). When fast gamma oscillations occur in the hippocampal CA1, the MEC also exhibits fast gamma oscillations. The oscillations from these two brain regions are highly synchronized, suggesting that fast gamma oscillations support signal transfer from the MEC to CA1. By contrast, synchronization of slow gamma oscillations suggests signal transfer from CA3 to CA1 (Colgin et al., 2009). We thus analyzed gamma oscillations in CA1 of WT and APP-KI mice. In WT mice, we observed gamma episodes across the slow and fast gamma range (20 – 90 Hz; Fig. 3B). However, in APP-KI mice, gamma oscillations were observed only in low frequency ranges, and marginal gamma episodes were observed at a frequency range above 40 Hz. Comparison of oscillatory power normalized by gross 1-100 Hz frequency power (Deshmukh et al., 2010) showed reduction of power at 62 – 100 Hz and 34 – 36 Hz ranges, and increase of power at 5 – 8 Hz theta oscillation range in APP-KI mice (false discovery rate (FDR) corrected for 100 multiple comparisons at 1–100 Hz bins, q<0.05; Fig. 3C).

**Figure 3.**
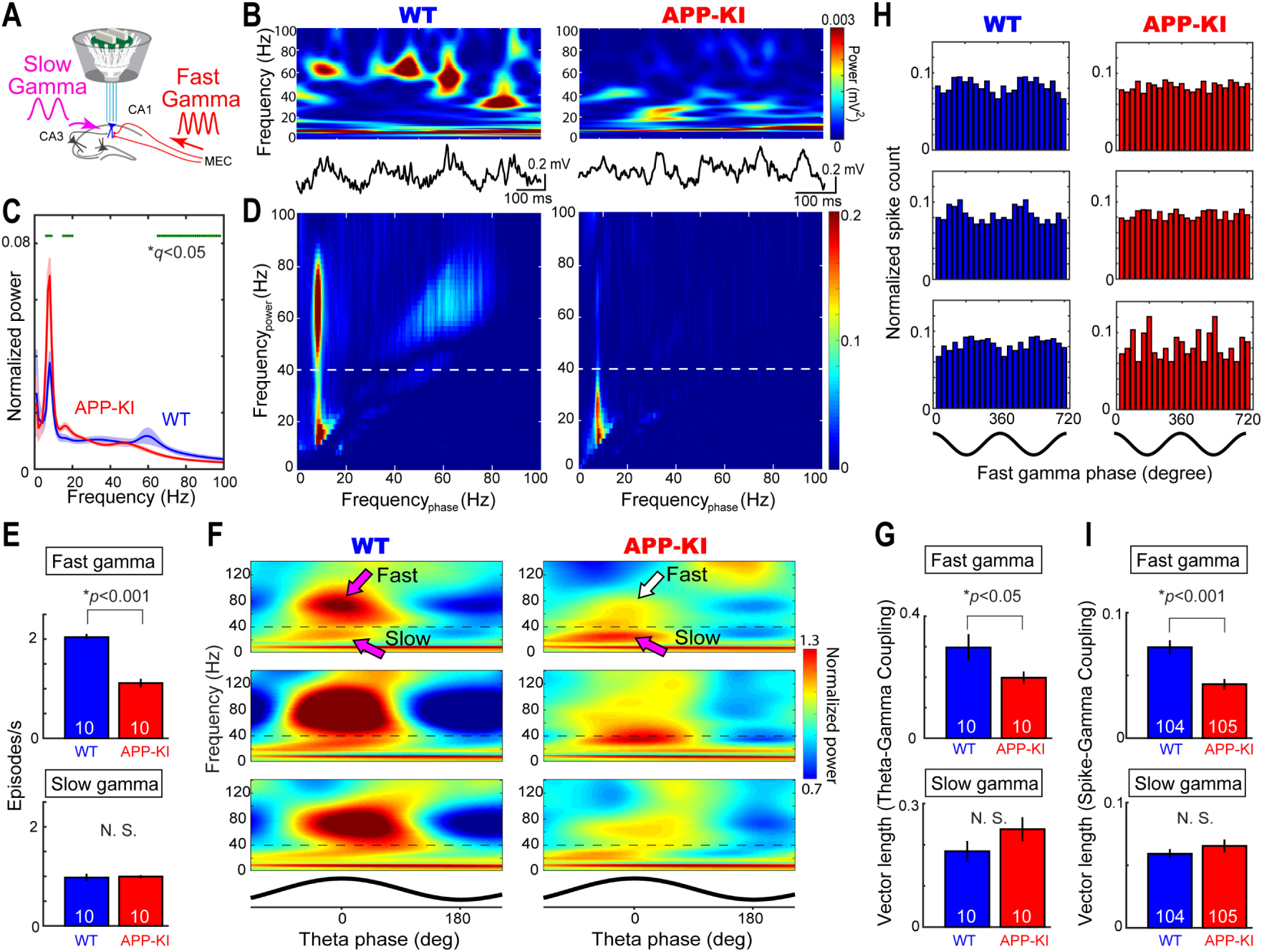
Fast gamma oscillations, but not slow gamma oscillations, were diminished in the hippocampal CA1 of APP-KI mice. (**A**) Fast and slow gamma oscillations that reflect inputs from MEC and CA3, respectively, were recorded in the CA1 during the open field task (see text). (**B**) A representative time-resolved spectrogram showing gamma oscillation episodes in a WT mouse (left) and APP-KI mouse (right). Corresponding raw local field potential traces are shown at the bottom. (**C**) Normalized power spectra from WT and APP-KI mice (n=10 mice each). Significant difference in power for each 1Hz bin is shown with green dot at the top (q<0.05 at ranges of 5-8 Hz, 34-36 Hz and 62-100 Hz, FDR corrected for multiple comparison). (**D**) Example cross-frequency coherence plots showing that oscillatory powers at 20–40 Hz and 40–80 Hz frequency (y-axis) were modulated by theta phase (x-axis) in a WT mouse (left), but the theta modulation of 40–80 Hz oscillatory power was absent in an APP-KI mouse (right). Dotted line denotes 40-Hz border between fast and slow gamma. (**E**) Occurrence rates of fast gamma episodes (top) and slow gamma episodes (bottom) for WT (n=10) and APP-KI mice (n=10). p<0.001, Wilcoxon ranks sum test. (**F**) Normalized power spectrograms averaged across all theta cycles in three representative WT mice (left) and APP-KI mice (right). Theta phase is shown at the bottom. Dotted line denotes 40-Hz border between fast and slow gammas. (**G**) Mean vector length for the distribution of fast gamma episodes (top) and slow gamma episodes (bottom) across theta phase. p<0.05, Wilcoxon ranks sum test. (**H**) Fast gamma phase distribution of spikes from example three CA1 neurons from WT mice (left) and APP-KI mice (right). Schematics of two cycles of gamma waveform are shown in the bottom. (**I**) Mean vector length for the distribution of spikes across fast gamma phase (top) and slow gamma phase (bottom) for WT CA1 neurons (n=104 cells) and APP-KI CA1 neurons (n=105 cells). p<0.05, Wilcoxon rank sum test.

To test whether fast and slow gamma oscillations are deteriorated in CA1, we investigated temporal structures of gamma oscillations. Previous healthy animal studies showed that gamma oscillations occur mainly at the peak to falling phase of co-existing theta oscillations, a property termed theta-gamma coupling (Bragin et al., 1995; Colgin et al., 2009). The theta-gamma cross-frequency coupling correlates with animals’ memory performance and is suggested to be a mechanism for local information processing during memory demands (Tort et al., 2009; Axmacher et al., 2010). Cross-frequency coherence plots from WT mice revealed peaks at two gamma frequency ranges at 20 – 40 Hz and 40 – 90 Hz that were coupled to specific phases of theta oscillations (Fig. 3D). We thus defined slow gamma oscillations at 20 – 40 Hz frequency range, and fast gamma oscillations at 40 – 90 Hz range in our experiment. In APP-KI mice, only the slow gamma oscillations from 20 – 40 Hz range were coherent to co-existing theta oscillations (Fig. 3D). The occurrence rate of fast gamma oscillation was diminished in APP-KI mice (p<0.001, Wilcoxon rank sum test, Fig. 3E), whereas the occurrence rate of slow gamma remained similar (p >0.05, Wilcoxon rank sum test, Fig. 3E). Cross-frequency coupling spectrogram revealed fast and slow gamma oscillations in the peak to falling phase of theta oscillations in WT mice, whereas fast gamma oscillations were significantly lost in APP-KI mice (Fig. 3F). The degree of theta-gamma cross-frequency coupling was assessed by measuring the phase-locking vector length (Colgin et al., 2009). The analysis revealed diminished theta-fast gamma coupling, while theta-slow gamma coupling was spared (p<0.05 for fast gamma; p>0.05 for slow gamma, Wilcoxon ranks sum test; Fig. 3G). To test whether spike timing is entrained by fast and slow gamma oscillations, we examined the temporal organization of spike activities during slow and fast gamma oscillations (Colgin et al., 2009). Phase-locking of spikes to fast gamma oscillations was diminished in APP-KI mice (p<0.001, Wilcoxon rank sum test; Fig. 3H-I). Altogether, our results demonstrate that fast gamma oscillations were disrupted in the hippocampal CA1 of APP-KI mice, while slow gamma oscillations were relatively spared. These results suggest that MEC→CA1 signal transfer via fast gamma oscillations is deteriorated in APP-KI mice, whereas slow gamma-mediated CA3→CA1 signal transfer remains relatively intact.

### MEC neurons in APP-KI mice showed severe impairment in spatial tuning

Disrupted fast gamma oscillations observed in hippocampal CA1 suggest that neural activity in the MEC is deteriorated in APP-KI mice. To test this possibility, we next investigated spatial representation in the MEC. We implanted a recording device into the MEC, and performed unbiased recording of principal neurons in layers 2 and 3 of the dorsal MEC (n=61 neurons from n = 13 WT mice; n= 65 neurons from n = 13 APP-KI mice; Fig. 4A-C **and** Fig. S12-S13; no difference in the distribution of sampling positions between WT and APP-KI mice, Fig. S13). To test for their spatial tuning, MEC neurons were recorded while animals were randomly foraging through the 1×1 m open field task. Among the recorded MEC neurons from WT mice, a portion of cells exhibited spatially tuned multiple firing fields as grid cells (Fig. 4D). By contrast, most of the recorded MEC neurons in APP-KI mice showed scattered firing patterns around the open field (Fig. 4E). To evaluate their grid cell property, we analyzed the *gridness score* by computing the autocorrelation of their firing rate map (Sargolini et al., 2006) (Fig. 4D-F). A cumulative distribution curve of the gridness score for all recorded MEC neurons indicates that, compared to WT mice, APP-KI mice significantly lost a group of neurons with gridness score of more than ∼0.4 (p<0.05, KS test; Fig. 4F). To define grid cells, we used a cut-off of gridness score at 0.41, which was the 95th percentile of the distribution of gridness score based on the shuffled WT data (Langston et al., 2010) (Fig. 4F **and** S14). We found that, among recorded MEC neurons, 20% were grid cells in WT mice, whereas only 2% were grid cells in APP-KI mice (p<0.001, binomial test; Fig. 4F-G). The scattered firing of APP-KI neurons resulted in significantly reduced spatial tuning (p< 0.001, Wilcoxon rank sum test) and increased firing field area (p <0.01, Wilcoxon rank sum test, Fig. 4G). However, we did not observe any difference in their mean or peak firing rates, suggesting that MEC neurons in APP-KI mice did not show hyperactivity reported previously in severe transgenic AD models (Busche et al., 2008) (Fig. 4G **and** S14). The distribution of spatial information score for MEC neurons indicate that APP-KI mice lost a significant portion of neurons with high spatial information score, suggesting that aperiodic spatially-tuned MEC neurons (Miao et al., 2017) were also impaired in APP-KI mice (Fig. 4H **and** S15). These data indicate that spatial tuning of MEC neurons, including grid cells, was severely disrupted in APP-KI mice.

**Figure 4.**
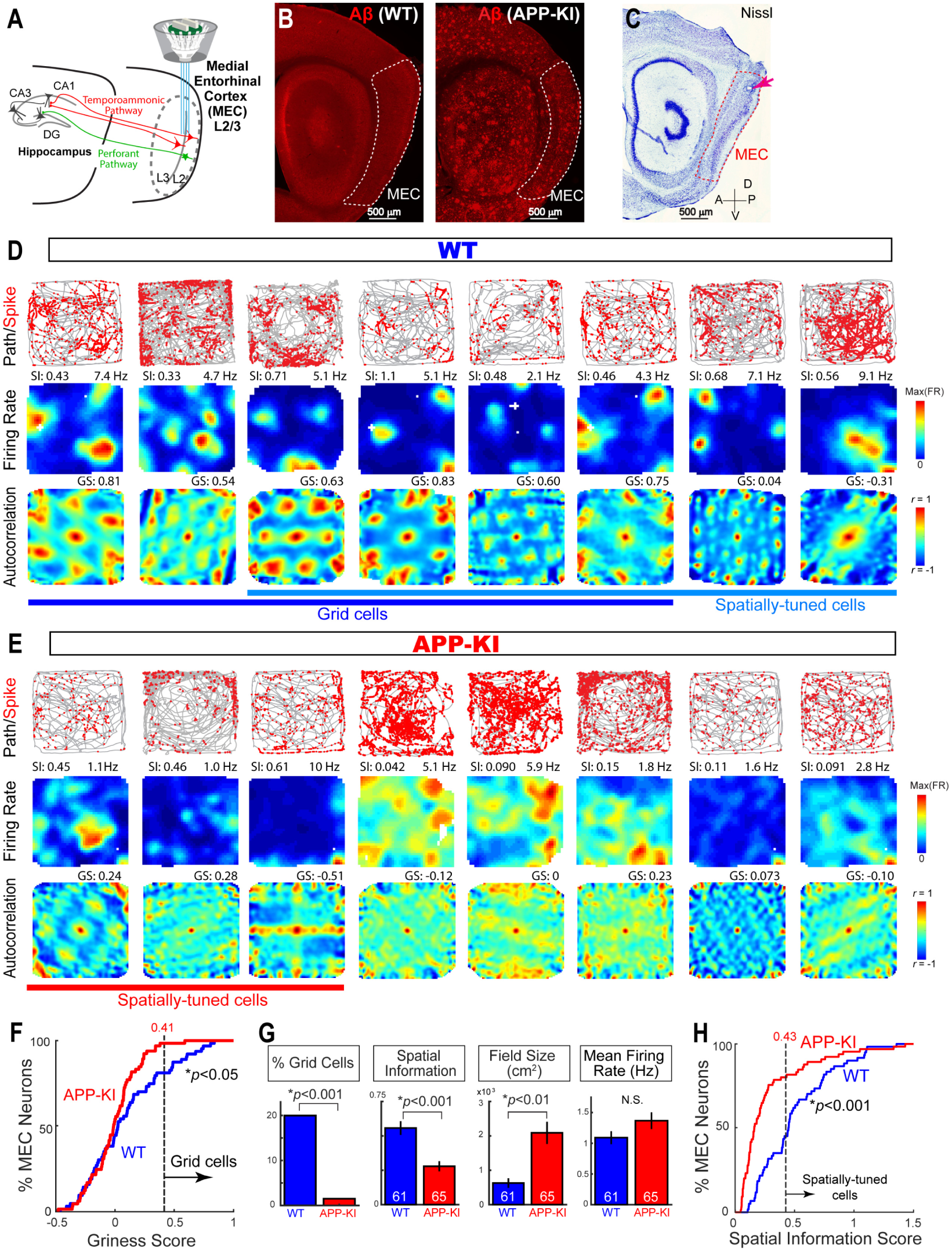
MEC neurons, including grid cells, showed impaired spatial tuning in APP-KI mice. (**A**) A 64-channel recording device was targeted to the layers 2/3 of the dorsal part of medial entorhinal cortex (MEC). (**B**) Sagittal sections with anti-Aβ immunostaining for 12-mo WT mouse (left) and 12-mo APP-KI mouse (right). (**C**) Sagittal sections with Nissl staining showing a representative recording position in the dorsal MEC (right, red arrow). Dotted lines denote the MEC. A, anterior; P, posterior; D, dorsal; V, ventral. Scale bars, 500 µm. (**D**) Eight representative MEC neurons from WT mice recorded in the 1×1 m open field. Top: Red dots and gray lines denote spike position and animal trace, respectively. Middle: Firing rate map. Color is scaled with maximum firing rate (Hz) shown at top right of each rate map. Spatial information (SI) score is shown at top left. Bottom: Autocorrelograms of the firing rate maps shown for 2×2 m range. Color is scaled with correlation from *r* = −1 to 1. Gridness score (GS) is shown at top right. (**E**) Eight representative MEC neurons as in (D), but from APP-KI mice. (**F**) Cumulative distribution plot for gridness score of MEC neurons in WT mice (blue, n=61) and APP-KI mice (red, n=65). A threshold of 0.41, obtained from 95^th^ percentile of shuffled WT data, was used to define grid cells. p<0.05, KS test. (**G**) From left to right: Percentage of neurons defined as grid cells (p<0.001, binomial test), spatial information score (p<0.001, Wilcoxon rank sum test), firing field size (p<0.01, Wilcoxon rank sum test), and mean firing rate (p>0.05, Wilcoxon rank sum test). (**H**) Cumulative distribution plots of spatial information score from all recorded MEC neurons from WT mice and APP-KI mice. A threshold of 0.43, obtained from 95^th^ percentile of shuffled WT data, was used to define spatially-tuned cells. p <0.001, KS test.

### MEC neurons in APP-KI mice could not send remapping signals to the hippocampus

We then tested whether remapping signals are properly signaled by these impaired MEC neurons. A previous healthy animal study showed that MEC grid cells exhibit remapping across distinct environments (Fyhn et al., 2007). The remapping of MEC grid cells occurs coherently with the remapping of hippocampal place cells, suggesting that MEC grid cells feed remapping signals to hippocampal place cells. Consistent with the previous finding, we observed that MEC neurons in WT mice showed distinct grid fields between Track A and Track B (Fig. 5A). However, MEC neurons in APP-KI mice showed scattered firing along both Track A and Track B, showing marginal difference in the firing patterns between the two tracks (Fig. 5B). PV correlation analysis revealed that APP-KI mice show larger PV correlations than that from WT mice, indicating that remapping property of MEC neurons is disrupted in APP-KI mice (p<0.001, KS text; Fig. 5C-D **and** S16-S17). We also assessed spatial correlation of individual neurons between Track A and Track B, confirming that MEC neurons showed disrupted remapping in APP-KI (Fig. 5E **and** S18). However, we did not observe significant difference between WT and APP-KI mice in the stability of MEC neurons from two Track A recordings (p > 0.05, KS test; Fig. 5C **and** 5E). These data demonstrate that remapping signals encoded by MEC neurons are severely impaired in APP-KI mice.

**Figure 5.**
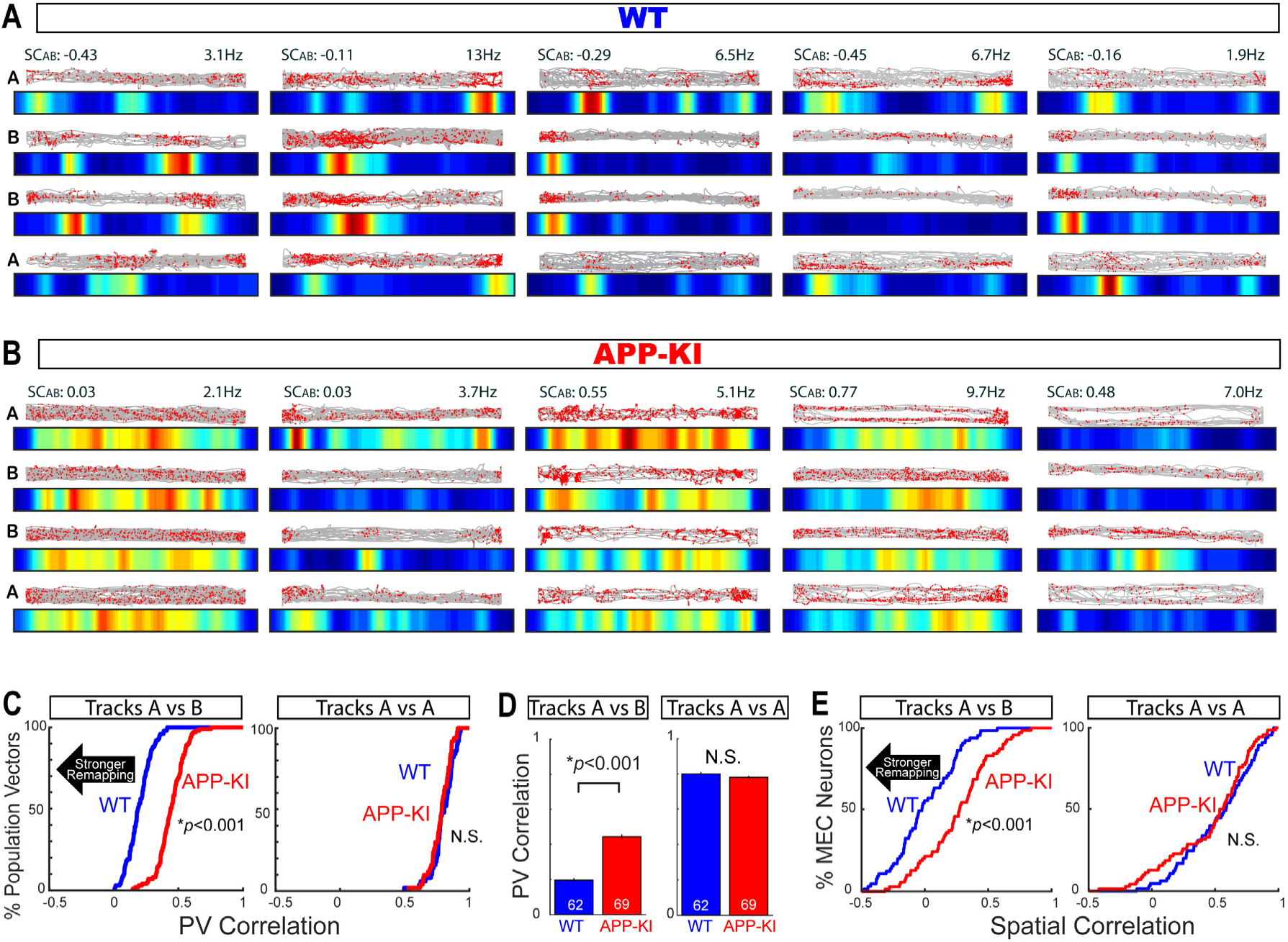
MEC neurons in APP-KI mice showed severely disrupted remapping. (**A**) Five representative MEC neurons from WT mice recorded in Tracks A, B, B, and A (from Top to Bottom). Red dots and gray lines at the top of each track denote spike position and animal trace in the 1 m linear tracks, respectively. Bottom color map denotes firing rate map. Color is scaled with maximum firing rate (Hz) shown at top right. Spatial correlation between Track A and B (SCAB) is shown at top left. (**B**) Five representative MEC neurons from APP-KI mice are shown as in (A). (**C**) Left: Cumulative distribution plots for PV correlation between Track A and Track B. Lower PV correlation denotes stronger remapping (black arrow)p<0.001, KS test. Right: Cumulative distribution plots for PV correlation between two recordings in Track A. p>0.05, KS test. (**D**) Left: Mean PV correlation between Track A and Track B. APP-KI mice showed significant increase (that is, weaker remapping) compared to that of WT mice. p<0.001, Wilcoxon rank sum test. Right: Mean PV correlation between two recordings in Track A did not differ. p>0.05, Wilcoxon rank sum test. (**E**) Left: Cumulative distribution plots of spatial correlation between Track A and Track B from MEC neurons during remapping task. Significant increase in APP-KI mice compared to that of WT mice indicates disrupted remapping. p<0.001, KS test. Right: Cumulative distribution plot of spatial correlation between two recordings in Track A. p >0.05, KS test.

### Remapping impairment co-existed with the disrupted spatial tuning in MEC neurons, but not in CA1 neurons

To examine the causality between impairment of spatial tuning and disruption of remapping in CA1 and MEC neurons, we measured the correlation between spatial information score and spatial correlation between Track A and Track B in individual neurons (Fig. 6). In WT mice, neither CA1 neurons nor MEC neurons showed correlation between spatial tuning and remapping, suggesting that these two functional properties are independent in healthy WT mice (p>0.05, Pearson correlation, Fig. 6A-B). However, in APP-KI mice, we found that MEC neurons show significant negative correlation between spatial information score and spatial correlation (p<0.001 and *r*(65) = −0.45, Pearson correlation, Fig. 6B). These results suggest that the impairment of spatial tuning causes remapping disruption in individual MEC neurons in a cell autonomous manner. By contrast, CA1 neurons in APP-KI mice did not show significant correlation (p>0.05, Pearson correlation, Fig. 6A), suggesting that the remapping disruption of individual CA1 neurons is not autonomously caused by the impairment of spatial tuning.

**Figure 6.**
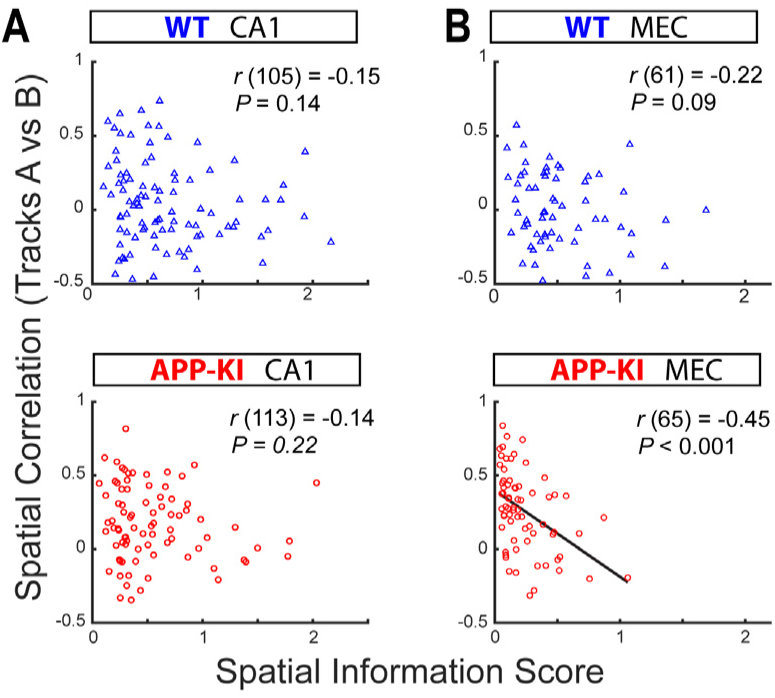
Remapping impairment co-existed with the disrupted spatial tuning in MEC neurons, but not in CA1 neurons of APP-KI mice. (**A**) Correlation between spatial tuning (spatial information score, x-axis) and remapping (spatial correlation for Tracks A vs Track B, y-axis). Each dot represents recorded CA1 neurons in WT mice (top) and APP-KI mice (bottom). (**B**) Correlation between spatial tuning and remapping as in (A), but for MEC neurons in WT mice (top) and APP-KI mice (bottom). Only MEC neurons in APP-KI mice showed significant correlation (p<0.001, *r* (65) = −0.45, Pearson correlation).

### Fast gamma coupling between CA1 and MEC were impaired in APP-KI mice

To test if the impairment of MEC neurons is accompanied by the deterioration of local gamma oscillations, we next examined gamma oscillations in the MEC (Fig. 7A). Because previous healthy animal studies found only fast gamma but not slow gamma oscillations in the MEC (Chrobak and Buzsaki, 1998; Colgin et al., 2009), we focused our analysis on fast gamma oscillations (no difference was observed for MEC slow gamma oscillations between WT and APP-KI mice, Fig. S19). The analysis of theta-fast gamma cross-frequency coupling revealed the absence of fast gamma oscillations coupled to theta oscillations (Fig. 7B). However, we did not see any difference in the normalized oscillatory power nor in the occurrence rate of fast gamma oscillation between WT and APP-KI mice (q>0.05 for normalized power, FDR corrected, Fig. 7C; p>0.05 for gamma occurrence, Wilcoxon rank sum test; Fig. 7D). Theta-phase spectrogram showed tendency of fast gamma oscillations being weakly modulated by theta oscillations (p=0.051, Wilcoxon rank sum test, Fig. 7E-F). The spikes from MEC neurons in WT mice were phase-locked to local fast gamma oscillations, as reported in previous study (Quilichini et al., 2010) (Fig. 7G-H). However, the spike-fast gamma phase-locking was markedly disrupted in APP-KI mice (p<0.01, Wilcoxon rank sum test, Fig. 7H **and** S20).

**Figure 7.**
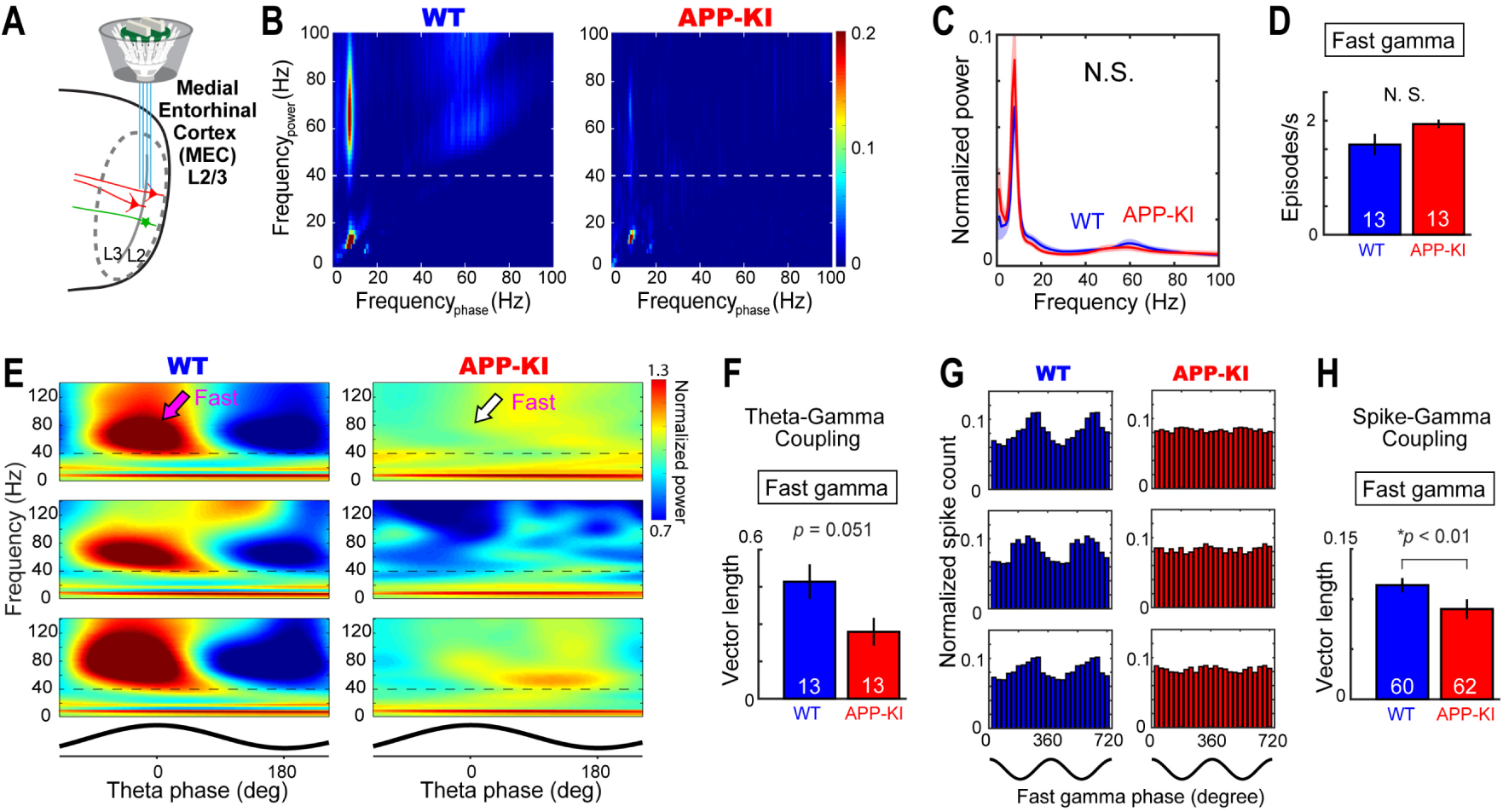
Impaired fast gamma oscillations in the MEC of APP-KI mice. (**A**) Fast gamma oscillations were recorded from the MEC during the open field task. (**B**) Example cross-frequency coherence plots showing that 40–100 Hz oscillatory power (y-axis) was modulated by theta phase (x-axis) in a WT mouse (left), but this modulation was absent in an APP-KI mouse (right). Dotted line denotes 40-Hz border between fast and slow gamma. (**C**) Normalized power spectra for WT and APP-KI mice (n=13 mice each). No difference was observed in the power spectra (q>0.05, FDR corrected for multiple comparison). (**D**) Occurrence rates of fast gamma episodes in WT and APP-KI mice. (**E**) Normalized power spectrograms averaged across all theta cycles in three representative WT mice (left) and APP-KI mice (right). Theta phase is shown at the bottom. Dotted line denotes 40-Hz border between fast and slow gammas. (**F**) Mean vector length for the distribution of fast gamma episodes across theta phase. p=0.051, Wilcoxon ranks sum test. (**G**) Fast gamma phase distribution of spikes from example three MEC neurons in WT (left) and APP-KI (right) mice. Schematics of two cycles of gamma waveform are shown in the bottom. (**H**) Mean vector length for the distribution of spikes across fast gamma phase for WT CA1 neurons (n=60 cells) and APP-KI CA1 neurons (n=62 cells). p<0.01, Wilcoxon ranks sum test.

To directly test the idea that the impairment of fast gamma oscillations affects MEC→CA1 signal transfer in APP-KI mice, we implanted two bundles of electrodes and simultaneously recorded from the hippocampal CA1 and the MEC (n = 6 WT mice and n = 6 APP-KI mice) (Fig. 8A). To test the degree of synchronization between the gamma oscillations in the MEC and the CA1, we assessed coherence of oscillatory activity (Igarashi et al., 2014). Analysis revealed that the gamma oscillations between the MEC and CA1 have weaker coherence at 58 – 75 Hz range in APP-KI mice (q<0.05, FDR corrected; Fig. 8B). We then collected oscillatory activity in the MEC when fast gamma oscillations were detected from the hippocampal CA1 (Colgin et al., 2009) (Fig. 8C). The MEC fast gamma oscillations were significantly desynchronized from the CA1 fast gamma oscillations in APP-KI mice. This was confirmed by the decreased fast gamma power ratio in APP-KI mice (p<0.05, Wilcoxon rank sum test; Fig. 8D). Altogether, these results demonstrate that the temporal coordination of the gamma oscillations between the MEC and the CA1 is diminished, suggesting that MEC→CA1 signal transfer mediated by the fast gamma oscillations is deteriorated in APP-KI mice.

**Figure 8.**
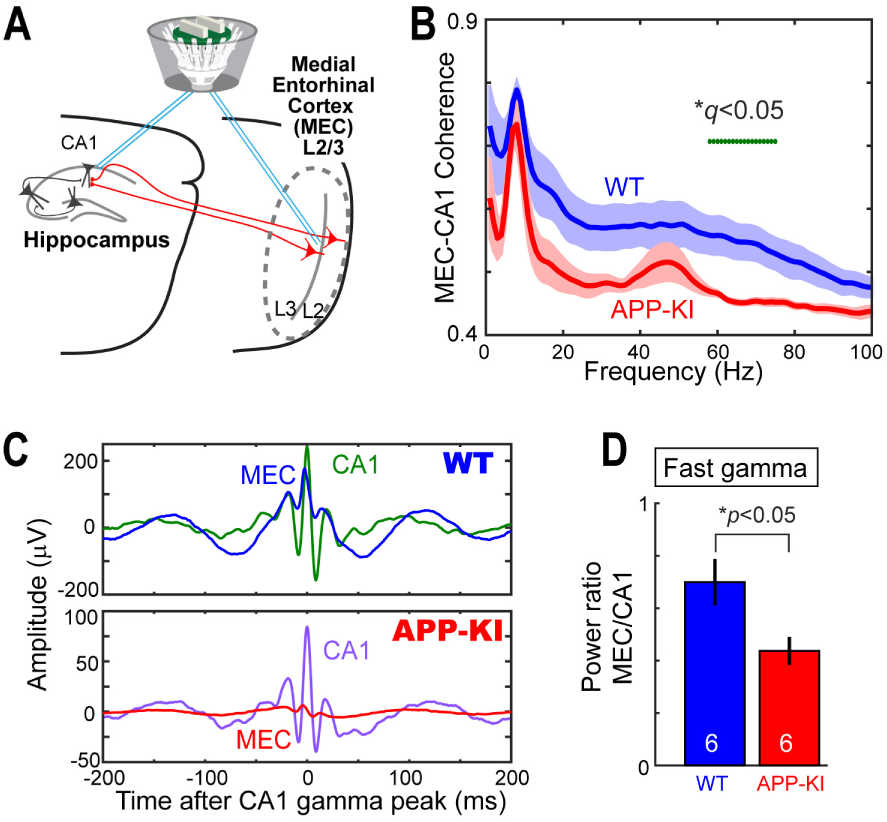
Disrupted coupling of fast gamma oscillations between the MEC and CA1. (**A**) Fast gamma oscillations were recorded simultaneously in the MEC and hippocampal CA1 during the open field task. (**B**) Mean coherence plot of oscillatory activity as a function of frequency in WT and APP-KI mice (n=6 mice each). Significant difference in coherence for each 1Hz bin is shown with green dot at the top (q<0.05 at 58-75 Hz range, FDR corrected for multiple comparison). (**C**) Representative averaged local field potential traces from the MEC of WT mouse (top) and APP-KI mouse (bottom) during fast gamma episodes detected in CA1. *t* = 0 corresponds to the peak amplitudes of the fast gamma episodes in CA1. (**D**) MEC:CA1 power ratio during fast gamma episodes detected in CA1 (p<0.05, Wilcoxon rank sum test).

## Discussion

In this study, using novel APP-KI mice, we investigated the effect of mutated APP on the remapping of MEC neurons and CA1 neurons in AD for the first time. We found that the remapping of CA1 neurons was disrupted in post-plaque APP-KI mice. Spatial tuning of MEC neurons, including grid cells, was severely impaired, sending disrupted remapping signals to the hippocampus. Fast gamma oscillations were deteriorated in both CA1 and MEC, suggesting impaired signal transfer in the MEC→CA1 circuit. This line of evidence points to the impairment of MEC neurons as an element of cause for remapping disruption in the entorhinal-hippocampal circuit of APP-KI mice (Fig. S1). Previous anatomical studies in AD patients showed that neurons in the entorhinal cortex exhibit the earliest neuronal deterioration (Van Hoesen et al., 1991; Gomez-Isla et al., 1996), but the link between deterioration of entorhinal neurons and spatial memory impairment has been unclear. Although impairments in the spatial tuning of place cells or grid cells were reported (Cacucci et al., 2008; Kunz et al., 2015; Fu et al., 2017; Mably et al., 2017), no previous studies have investigated remapping of place cells and grid cells in AD subjects. Our study provides the first circuit-level evidence that identifies the link between the impairment of MEC neurons and the remapping disruption of hippocampal CA1 neurons. In early-stage AD patients, deterioration of entorhinal neurons may trigger remapping impairment in hippocampal neurons. Because the remapping of hippocampal neurons is one of the most plausible circuit functions for spatial pattern separation to distinguish multiple environments (Colgin et al., 2008; Yassa and Stark, 2011), disruption of remapping would ultimately lead to spatial memory loss and wandering symptom in AD. Although the hippocampus has been used as the main model brain region in AD research for decades, our finding strengthens the importance of entorhinal cortex as a potential area of disease origin. Together with a previous study showing diminished grid cell activity in humans at AD risk (Kunz et al., 2015), our results also point to entorhinal cortex as a potential target for future diagnostic and therapeutic endeavor against AD.

In healthy animals, neurons in the entorhinal-hippocampal circuit exhibit two forms of remapping known as “global remapping” and “rate remapping” (Leutgeb et al., 2005b). When animals move across environments with distinct features, entorhinal grid cells and hippocampal place cells show distinct patterns of firing (global remapping). Global remapping of place cells and grid cells occur simultaneously (Fyhn et al., 2007), and inhibition of entorhinal grid cells scrambles remapping of hippocampal place cells (Miao et al., 2015). These findings suggest that MEC input plays a crucial role in global remapping of hippocampal place cells. By contrast, when there is only a slight change in an environment, hippocampal place cells show change in their firing rate, while keeping the position of firing field constant (rate remapping). Because grid cells do not exhibit rate remapping (Fyhn et al., 2007), and because neurons in the hippocampal CA3 and dentate gyrus exhibit the most pronounced rate remapping (Leutgeb et al., 2005a; Leutgeb et al., 2007), rate remapping signal is suggested to be mainly generated inside the dentate gyrus/CA3. Although it is likely that both global remapping and rate remapping are needed for high-precision spatial memory, disruption of global remapping would completely deprive AD patients of spatial information about which environment they are currently in. Our results showing the disruption of global remapping in the MEC and CA1 of APP-KI mice clearly explain the devastating impairment of spatial memory in AD patients, who often exhibit wandering symptom and cannot discriminate distinct features of surrounding environments. Future development of therapeutic methods to protect or reactivate remapping in the entorhinal cortex may lead to a powerful tool that can be used to slow the rate of spatial memory decline in AD patients.

## Acknowledgments

We thank Jason Lee, Tomoaki Nakazono, Tomoko Viaclovsky, Kaori Shiraiwa, Anish Reddy, Allen Bramian and Ayushi Patel for technical assistance, and Mathew Blurton-Jones, Kim Green, Masashi Kitazawa, Robert Hunt and members of Igarashi lab for helpful discussion.

## Author contributions

K.M.I. and H.J. designed experiments and analyses; H.J. and K.M.I. performed the experiments; H.J. and K.M.I. performed the analyses, with input from S.S.; T.S and T.C.S generated the APP-KI mice; H.J. and K.M.I wrote the paper with input from all authors.

## Data and materials availability

APP-KI mouse is available at RIKEN Bio Resource Center (RBRC06344). Neurophysiological data and analytical codes are available upon request.

## Supplementary materials

### Materials and Methods

#### Subjects

Mice were maintained in standard housing conditions on a 12h dark/light cycle with food and water provided ad libitum. All procedures were conducted in accordance with the guidelines of the National Institutes of Health and approved by the Institutional Animal Care and Use Committee at the University of California, Irvine. We previously generated App^NL-G-F/NL-G-F^ maintained on a C57BL/6 background (Saito et al., 2014). For all experiments, eighteen App^NL-G-F/NL-G-F^ mice (fourteen males and four females) and seventeen App^WT/WT^ mice (eleven males and six females) between 7 months and 13 months of age were used. We noted no difference in our data between sexes. Detailed information about ages when the recording was performed is in Supplementary Table S1. Animals were housed in a reversed 12h dark/light cycle, and all testing occurred during the dark phase. If animals died or were sick during or before the final test, their recording positions could not be validated and therefore were removed from the data analysis.

#### Surgery and Electrode Preparation

All mice received a custom-built 64-channel drive modified from Liang et al (Liang et al., 2017) targeting either CA1 or MEC. For a group of animals, eight tetrodes targeted CA1 of hippocampus and another eight tetrodes targeted the superficial layer of medial entorhinal cortex. Tetrodes were constructed from four twisted 17 μm polyimide-coated platinum-iridium (90%–10%) wires (California Fine Wire). The electrode tips were plated with gold to reduce electrode impedances to between 150 and 300 kΩ at 1 kHz. Animals were anesthetized with isoflurane (air flow: 0.8–1.0 l/min, 1% isoflurane, adjusted according to physiological condition). The mice received subcutaneous injections of buprenorphine at the start of the surgery. Depth of anesthesia was examined by testing tail and pinch reflexes as well as breathing. Upon induction of anesthesia, the animal was fixed in a Kopf stereotaxic frame for implantation. A craniotomy was performed centering at AP 2.5 mm ML 2.5 mm from Bregma for hippocampus (approximately 1 x 1mm) and 0.2-0.4mm anterior to transverse sinus and ML 3.5mm from midline for medial entorhinal cortex (approximately 1 x 1mm). The dura was carefully removed, and the tetrodes were implanted. A stainless-steel screw fixed to skull above the cerebellum served as the ground electrode. The recording drive was secured to the scratched skull using dental cement.

#### Apparatus and Training Procedures

Behavioral training started at least 5 days after the surgery and data collection was performed exactly on the fifth day of the training. All experiments were conducted in a room containing a 1m-square box made of black acryl and a polarizing white cue card (216mm x 280mm) and two 1-m long linear tracks. In the open field task, running was motivated in the 1m-square box with cookie crumbs thrown randomly into the enclosure. Each session lasted 10 to 20 min. Upon finishing the open field task, animals rested for 5 minutes and then were tested in the remapping task. The remapping task training required mice to run in 1-m long linear tracks. Two 1-m long linear tracks were used: (Track A) Black acryl enclosure with inner wall decorated with patterns of white cue cards across the track and black rubber floor. (Track B) White acryl enclosure with inner wall decorated with distinct patterns of black cue signs across the track and white sandpaper floor. During the remapping task, animals were trained to run in successive series of Track A → Track B → Track B → Track A. Between the sessions, animals were given 5 minutes rest. Running was motivated by placing cookie crumbs on the end positions of the linear tracks. Each session lasted 5 to 10 min. On the linear tracks, the mice ran 10 full laps (back and forth).

#### Data Collection

LFPs were recorded single-ended using a ground in the skull above the cerebellum. One mouse without single-ended recording was excluded for LFP analyses. The tetrodes were connected to a multichannel, impedance matching, unity gain headstage (Neuralynx). The output of the headstage was conducted to a data acquisition system (Neuralynx). Unit activity was amplified by a factor of 3000–5000 and band-pass filtered from 600 Hz to 6000 Hz. Spike waveforms above a threshold set by the experimenter (∼55 μV) were time-stamped and digitized at 32 kHz for 1 ms. LFP signals, 1 per tetrode, were recorded in the 0–475 Hz frequency band at a sampling rate of 2000 Hz. Notch filters were not applied. The recording system tracked the position of two light-emitting diodes (LEDs), one large and one small, on the head stage (sampling rate 50 Hz) by means of an overhead video camera.

#### Data Analysis

Unless indicated otherwise, analyses were performed using MATLAB codes kindly provided by Dr. Edvard Moser, or written by the authors (Igarashi et al., 2014)

#### Spike Analysis

##### Spike Sorting and Cell Classification

Spike sorting was performed offline using graphical cluster-cutting software, MClust (A.D. Redish). Putative excitatory cells were distinguished from putative interneurons by spike peak-valley width and average rate. (Bartho et al., 2004). All putative excitatory cells with spike peak-valley width of more than 230 µs and mean firing rates of more than 0.1 Hz (mean firing rate during all left and right cue sampling intervals) were included for further analysis.

##### Firing rate maps

Position estimates were based on tracking of the LEDs on the head stage connected to the recording drive. For rate maps in the random foraging task, data were speed-filtered; only epochs with instantaneous running speeds of 1 cm/s or more were included.

To characterize firing fields, the position data were sorted into one-dimensional 1 cm bins for the linear tracks and 2.5 cm × 2.5 cm bins for the open field. For the open field map, the path was smoothed with a 21-sample boxcar window filter (400 ms; 10 samples on each side). Firing rate distributions were then determined by counting the number of spikes in each bin as well as the time spent per bin. Maps for number of spikes and time were smoothed individually using a boxcar average over the surrounding 5 × 5 bins. Weights were distributed as follows:

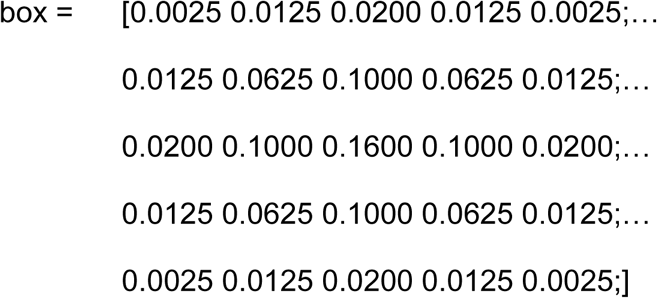

##### Spatial information score

For each cell, the spatial information score in bits per spike was calculated from the recordings in the open field task as

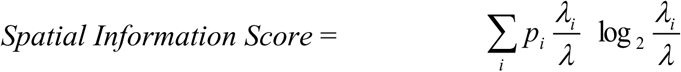

where *λ_i_* is the mean firing rate of a unit in the *i*th bin, *λ* is the overall mean firing rate, and *p_i_* is the probability of the animal being in the *i*th bin (occupancy in the *i*th bin / total recording time) (Skaggs et al., 1993). An adaptive smoothing method, introduced by Skaggs *et al.* (Skaggs et al., 1996), was used before the calculation of information scores (Henriksen et al., 2010).

##### Defining Place Cells

Place cells were defined as cells with spatial information score above the chance level (Langston et al., 2010). The chance level was determined in each brain region by a random permutation procedure using all cells recorded at that time-point in that region. One hundred permutations were performed for each cell in the sample. For each permutation trial, the entire sequence of spikes fired by the cell was time-shifted along the animal’s path by a random interval between 20 s and 20 s less than the total length of the trial (usually 600 −20 = 580 s), with the end of the trial wrapped to the beginning to allow for circular displacements. This procedure allowed the temporal firing structure to be retained in the shuffled data at the same time as the spatial structure was lost. Spatial information was then calculated for each shuffled map. The distribution of spatial information values across all 100 permutations of all cells in the sample was computed and finally the 95th percentile was determined. Place cells were defined as cells with spatial information scores above the 95th percentile of the distribution from shuffled data for the relevant group.

##### Gridness score

The structure of all rate maps was evaluated for all cells by calculating the spatial autocorrelation for each smoothed rate map (Sargolini et al., 2006; Fyhn et al., 2007). Autocorrelograms were based on Pearson’s product moment correlation coefficient with corrections for edge effects and unvisited locations. With *λ (x, y)* denoting the average rate of a cell at location *(x, y)*, the autocorrelation between the fields with spatial lags of *τ_x_* and *τ_y_* was estimated as:

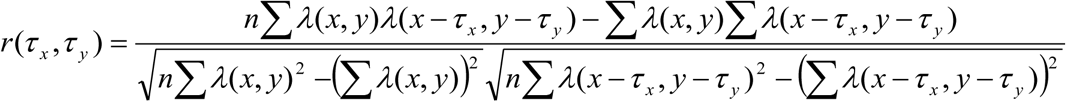

where the summation is over all n pixels in *λ (x, y)* for which rate was estimated for both *λ (x, y)* and *λ (x – τ_x_, y – τ_y_)*. Autocorrelations were not estimated for lags of *τ_x_, τ_y_* where *n* < 20.

The degree of spatial periodicity (‘gridness score’) was determined for each recorded cell by taking a circular sample of the autocorrelogram, centered on the central peak but with the central peak excluded, and comparing rotated versions of this sample. The Pearson correlation of this circle with its rotation in α degrees was obtained for angles of 60° and 120° on one side and 30°, 90° and 150° on the other. The celĺs gridness score was defined as the minimum difference between any of the elements in the first group and any of the elements in the second. The radius of the excluded central peak was defined as either the first local minimum in a curve showing correlation as a function of average distance from the center, or as the first incidence where the correlation was negative, whichever occurred first.

##### Defining Grid Cells

Grid cells were defined as cells in which rotational symmetry-based gridness scores exceeded the 95th percentile of a distribution of grid scores for shuffled recordings from the entire population of cells. Shuffling was performed in the same way as for place cells, i.e. for each permutation trial, the entire sequence of spikes fired by the cell was time-shifted along the animal’s path by a random interval between 20 s and 20 s less than the length of the trial (usually 600 – 20 = 580 s), with the end of the trial wrapped to the beginning. The shuffling procedure was repeated 100 times for each of the cells in the sample. For each permutation, a firing rate map and an autocorrelation map were constructed and a grid score was calculated. Grid scores were computed for all permutations for all cells and the 95th percentile was read out from the overall distribution. Grid cells were defined as cells with grid scores higher than the 95th percentile of the grid scores of the distribution for the shuffled data.

##### Remapping Analysis

Remapping analysis was performed as described previously (Miao et al., 2015). For experiments on the linear track, instantaneous firing rates on individual laps were estimated using a Gaussian kernel on the spike data for temporal smoothing:

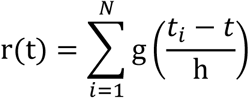

where g is a 1D Gaussian kernel, h is bandwidth, N is the total number of spikes, and t_i_ is the time of the i^th^ spike. The bandwidth was set at 100 ms. Due to minimal coverage per bin, temporal smoothing gives more robust rate estimates compared to those based on spatial bins during fast running on the track. Spike rate at each track position (1 cm bin) on each lap was estimated using a linear interpolation method applied to the temporally-smoothed spike rates. Because the same cells often had different firing fields on left and right runs, firing fields of the same cell in each run direction were analyzed as two distinct data sets. A segment of 5 cm was excluded from each end of the linear track for analysis.

Spatial correlation was obtained by calculating the Pearson correlation coefficient for mean firing rates across 1 cm wide bins from the track on a pair of sessions. For population vector analysis, population vectors were defined for each 3.6-cm bin of rate maps (25 bins in total) from all cells in the experimental group. To maintain independence of neighboring spatial bins, we estimated mean firing rates in spatial bins without smoothing. For lap-by-lap population vector analyses, a total of 10 laps (defined as pairs of forward and backward runs) were taken from each recording session. Population vector correlations were determined for each spatial bin of each lap.

#### LFP analysis

Animals with single-ended LFP recordings were used for the analysis.

##### Time–frequency analysis of power

Time-resolved power across frequencies was computed using a wavelet transform method. Signals were convolved by a family of complex Morlet’s wavelets *w*(*t*, *f*), one for each frequency, as a function of time:

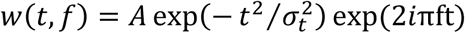

in which the family of wavelets was characterized by a constant ratio *f*/*σ_f_*, which was set to 7, and with *σ_f_* = ½π*σ_t_*. The coefficient *A* was set at:

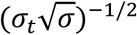

in order to normalize the wavelets such that their total energy was equal to 1.

##### Normalization of LFP power

To control for impedance differences between tetrodes, LFP power was normalized, for each tetrode, to the total power in 1–100 Hz band.

##### Bandpass filtering

An acausal (zero phase shift), frequency domain, finite impulse response bandpass filter was applied to the signals. For theta filtering, 5 and 6 Hz were chosen for the stopband and passband, respectively, for the low cut-off frequencies; 10 and 11 Hz were chosen for passband and stopband high cut-off frequencies. For filtering of gamma oscillations, 19 and 20 Hz were chosen for the stopband and passband, respectively, for the low cut-off frequencies; 90 and 91 Hz were chosen for passband and stopband high cut-off frequencies. For filtering of slow gamma oscillations, 19 and 20 Hz were chosen for the stopband and passband, respectively, for the low cut-off frequencies; 40 and 41 Hz were chosen for passband and stopband high cut-off frequencies. For filtering of fast gamma oscillations, 39 and 40 Hz were chosen for the stopband and passband, respectively, for the low cut-off frequencies; 90 and 91 Hz were chosen for passband and stopband high cut-off frequencies.

##### Selection of theta cycles

Theta cycles were selected by bandpass filtering the signal from 6-10 Hz and selecting local minima in the filtered signal (i.e., θ(t – 1) > θ(t) and θ(t + 1) > θ(t)). Segments of the recording were collected and defined as a theta cycle if the time between detected points fell within a range criterion that corresponded to the period of an ∼8 Hz theta cycle (i.e. 125 ± 25 ms). Local minima of detected theta cycles were required to be separated by at least 100 ms.

##### Detection of oscillation episodes

To extract periods of gamma oscillatory activity in the LFP, we first computed time-varying power within the frequency bands for each recording. Power at each time point was averaged across the frequency range to obtain time-varying estimates of oscillatory power. Time points were collected when the power exceeded 2 SD of the time-averaged power. Time windows, 160 ms in length, were cut around the identified time points. In each 160 ms segment, the maxima of gamma oscillatory amplitude were determined from the gamma bandpass filtered versions of the recordings. Duplicated gamma oscillatory periods, a common consequence of extracting overlapping time windows, were avoided by discarding identical maxima values within a given gamma oscillatory subtype and further requiring that maxima of a given subtype be separated by at least 100 ms. Individual gamma oscillatory windows were finally constructed from the original, non-bandpass filtered recordings as 400 ms long windows centered around the gamma oscillatory amplitude maxima.

##### Relationship of gamma to theta phase

LFP recordings were bandpass-filtered in the theta range (6–10 Hz), and theta phases for each time point were estimated using the Hilbert transform function from the Signal Processing Toolbox in MATLAB. Theta phases at the time points associated with gamma maxima (determined as described above) as well as theta phases for spikes were collected. Theta phases for each gamma oscillatory event were sorted into 30° bins, allowing the phase distribution of each event to be determined. For a given recording, the distributions of gamma oscillations were normalized by dividing the bins by the total number of gamma oscillatory episodes within a given recording. In this analysis, and in all analyses involving oscillation phase, the oscillation peak was defined as 0°.

##### Time–frequency representation of power across individual theta cycles

Time-varying power in 2-Hz-wide frequency bands, from 2 Hz to 140 Hz, was obtained for individual theta cycles using the wavelet transform method described above. Time frequency representations for multiple theta cycles recorded from the same site and session within the same animal were then averaged.

##### Strength of theta-gamma coupling

The theta phase at the time of gamma oscillation maxima occurrence was calculated by bandpass-filtering the LFP in the theta range, performing a Hilbert transform on the filtered signal, and then locating theta phase of individual gamma oscillation maxima. Resultant vectors were calculated from the phase distributions of gamma maxima. Lengths of resultant vectors were used as an index for the strength of theta-gamma coupling.

##### Phase-locking of spikes to gamma oscillations and theta oscillations

Neurons recorded in animals with single-ended LFP recordings were used for the analysis. The oscillatory phase at the time of spike occurrence was estimated. This was performed by first bandpass-filtering the LFP, performing a Hilbert transform on the filtered signal, and then extracting the phase component at the spike times. Cells were considered to be phase-locked if their phase distribution differed significantly from a uniform distribution (P < 0.05, Rayleigh test). Resultant vectors were calculated from the distributions of phase component. Lengths of resultant vectors were used as an index for the strength of phase-locking.

#### Histology and Reconstruction of Recording Positions

Electrode positions were confirmed by anaesthetizing the drive-implanted mice using isoflurane and performing small electrolytic lesions through passing current (10 μA for 20 s) through the electrodes. Immediately after this, the mice received an overdose of isoflurane and were perfused intracardially with saline followed by 4% freshly depolymerized paraformaldehyde in phosphate buffer (PFA). The brains were extracted and stored in the same fixative overnight. After overnight cryoprotection in phosphate buffer saline with 30% sucrose at 4°C, tissue samples were embedded in O.C.T. mounting medium and sagittal sections (40 μm) were cut and stained with cresyl violet. All tetrodes were identified, and the tip of each electrode was found by comparison with adjacent sections. Only data from tetrodes in the hippocampal CA1 or the layer 2/3 of dorsal MEC was collected for analysis.

For immunostaining, sections were rinsed 3 times for 10 min in 1 x PBS (pH 7.6) at room temperature, preincubated for 1 hr in 10% normal goat serum in PBST (1 x PBS with 0.5% Triton X-100). Between incubation steps, sections were rinsed in PBST. Sections were incubated with antibodies against Aβ, raised in mouse (MCSA1, Medimabs, 1:1000) for 24 hr in antibody-blocking buffer at 4°C. After three 15 min washes in PBST at room temperature, sections were incubated in a goat-anti mouse antibody conjugated with Alexa Fluor 568nm (Invitrogen, 1:250) for 2 hr at room temperature. After rinsing in PBS, sections were mounted onto glass slides with 40, 60 -diamidino-2-phenylindole (DAPI)-containing Vectashield mounting medium (Vector Laboratories), and a coverslip was applied. Digital photomicrographs were acquired with a Olympus BX53 fluorescent microscope equipped with a digital camera.

#### Statistical Analysis

All statistical testing assumed a non-parametric distribution and the Wilcoxon rank sum test was used. For comparing the distribution of place cells and grid cells, Kolmogorov-Smirnov (KS) tests were applied. All statistical methods used are summarized in Supplementary Tables S2-S14.

## Supplementary Figure Legends

**Fig. S1.**
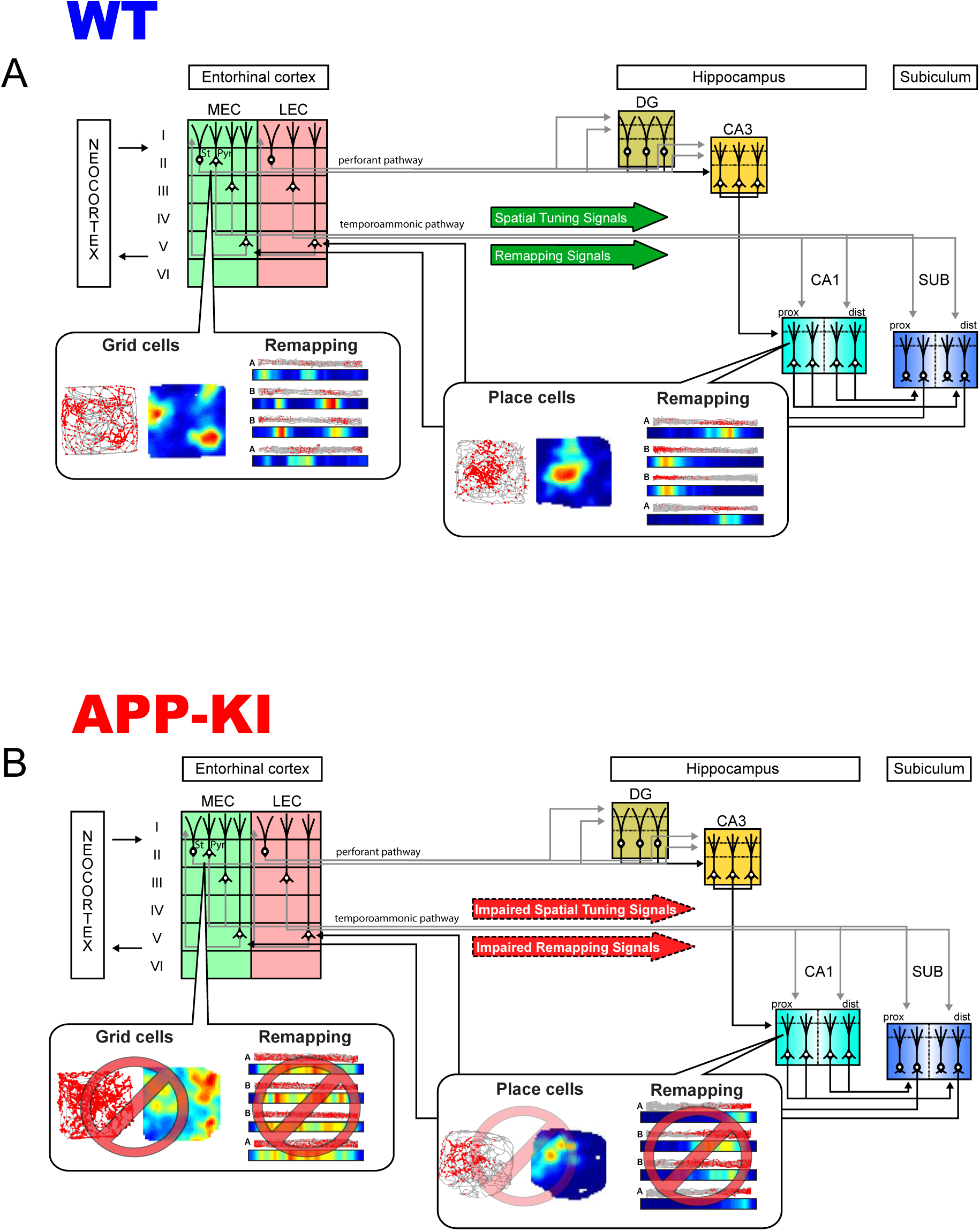
Schematics for parallel disruptions of entorhinal grid cells and hippocampal place cells. (**A**) In the entorhinal-hippocampal circuit of healthy brains, neurons in layers 2 and 3 of the medial entorhinal cortex (MEC) show spatially tuned neuronal activity and exhibit remapping across distinct environments. Pyramidal cells (Pyr) in layers 2 and 3 send these signals to neurons in the CA1 region of the hippocampus. In hippocampal CA1, neurons show spatial activity as place cells and exhibit remapping. Stellate cells (St) in MEC layer 2 send information to dentate gyrus and CA3. (**B**) In this study, we determined whether (1) remapping of hippocampal place cells is deteriorated in AD subjects, and if so, (2) whether place cell deterioration is linked to impairment of entorhinal grid cells, or independent from grid cells. We found that hippocampal CA1 neurons showed severely disrupted remapping, but mildly diminished spatial tuning in APP-KI mice. We also found that entorhinal grid cells were severely impaired, sending disrupted remapping signals to the hippocampus. These results point to the link between grid cell impairment and remapping disruption as an underlying circuit mechanism causing spatial memory loss in AD patients, who often cannot discriminate distinct features of surrounding environments.

**Fig. S2.**
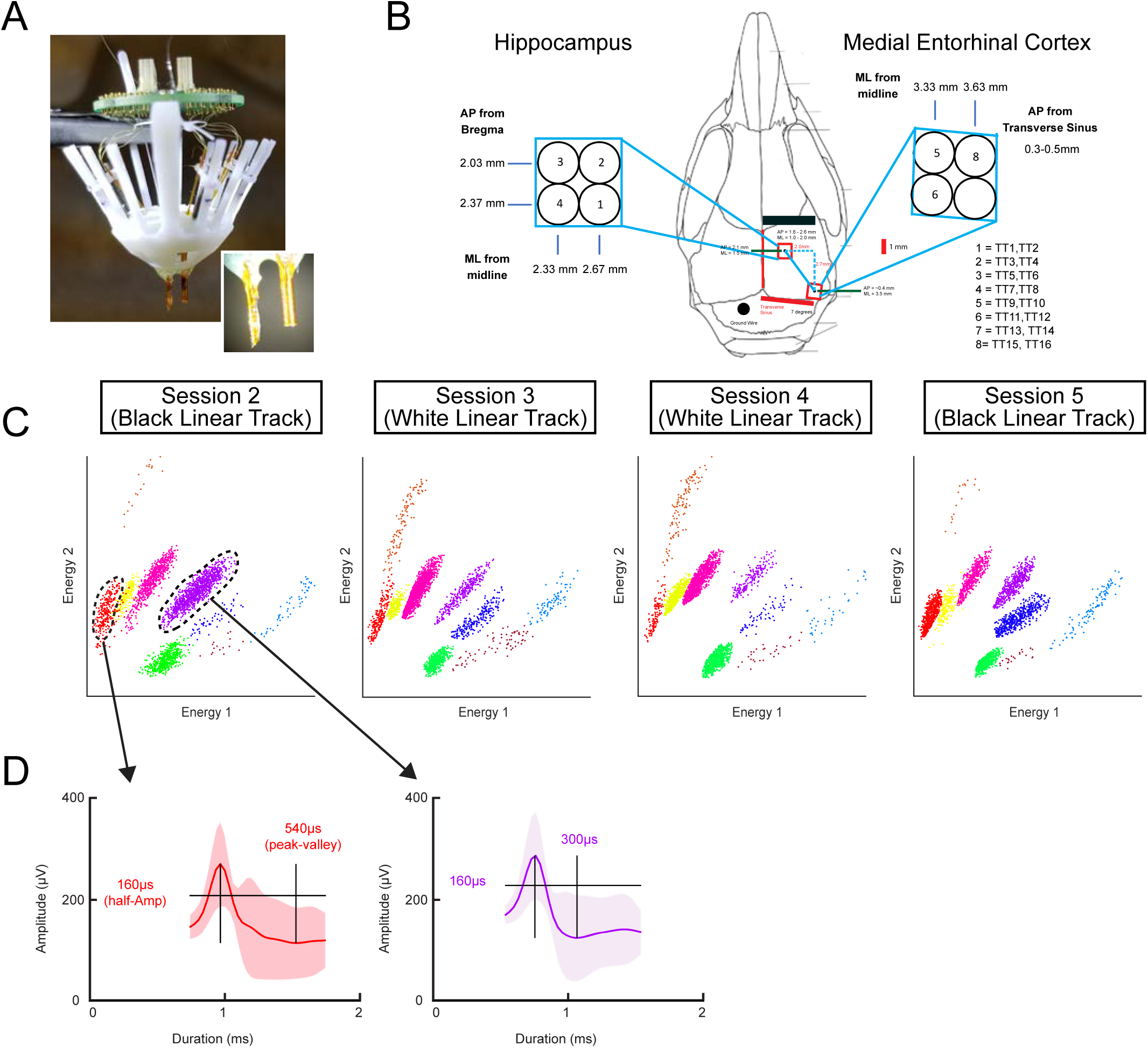
Electrophysiological recording of principal neurons in CA1 and MEC. (**A**) Multi-electrode 64-channel drives were used for CA1 and MEC neural recordings. Inset, magnification image showing two separate bundles containing tetrodes. (**B**) Surgical scheme illustrating the location of tetrodes implanted in CA1 of hippocampus and layer II/III of medial entorhinal cortex. (**C**) Example of spike sorting in energy space to identify single units across sessions, demonstrating stable chronic recording across sessions. (**D**) Two clusters of spikes and their corresponding spike wave forms from identified clusters representing two single units.

**Fig. S3.**
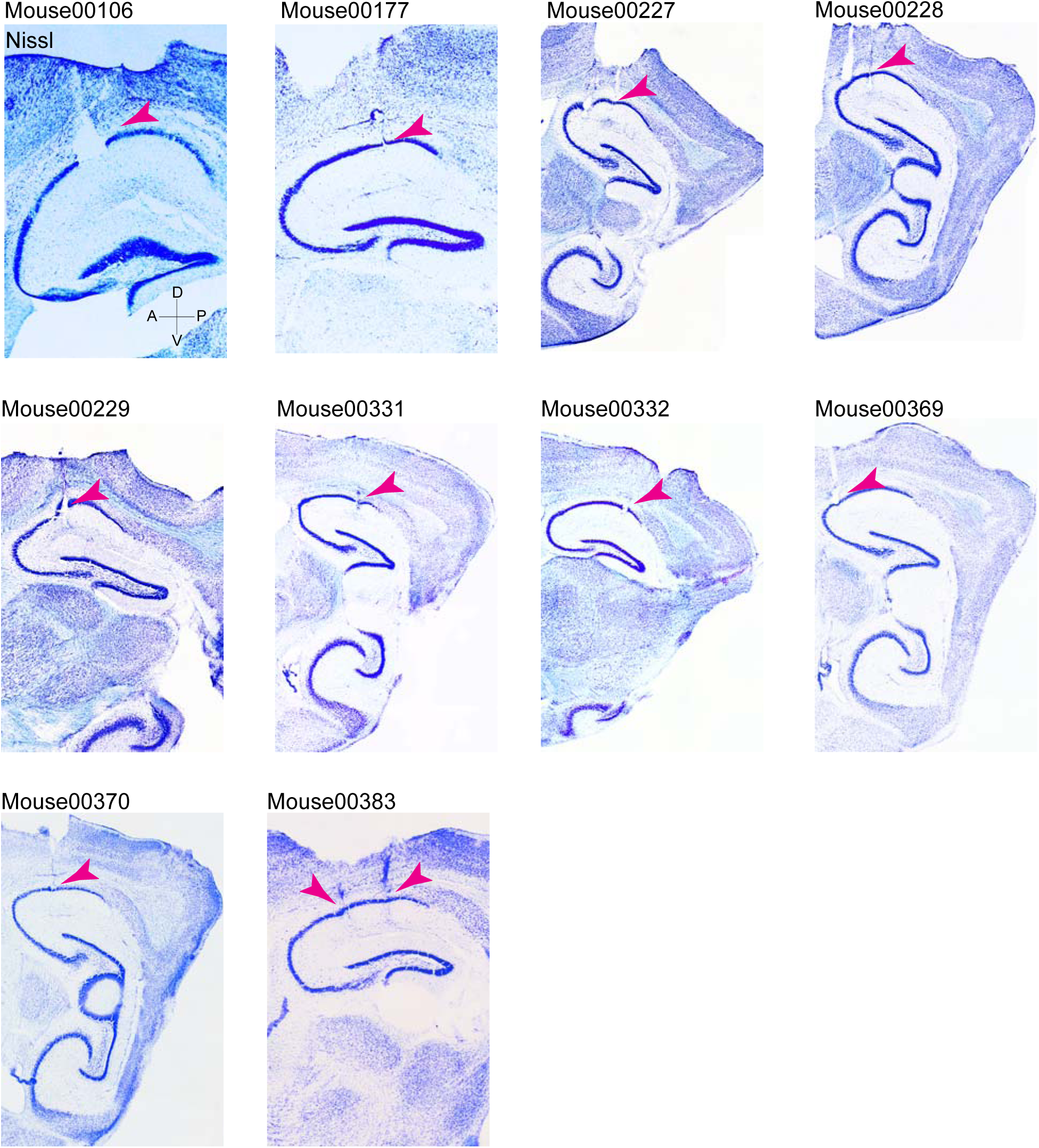
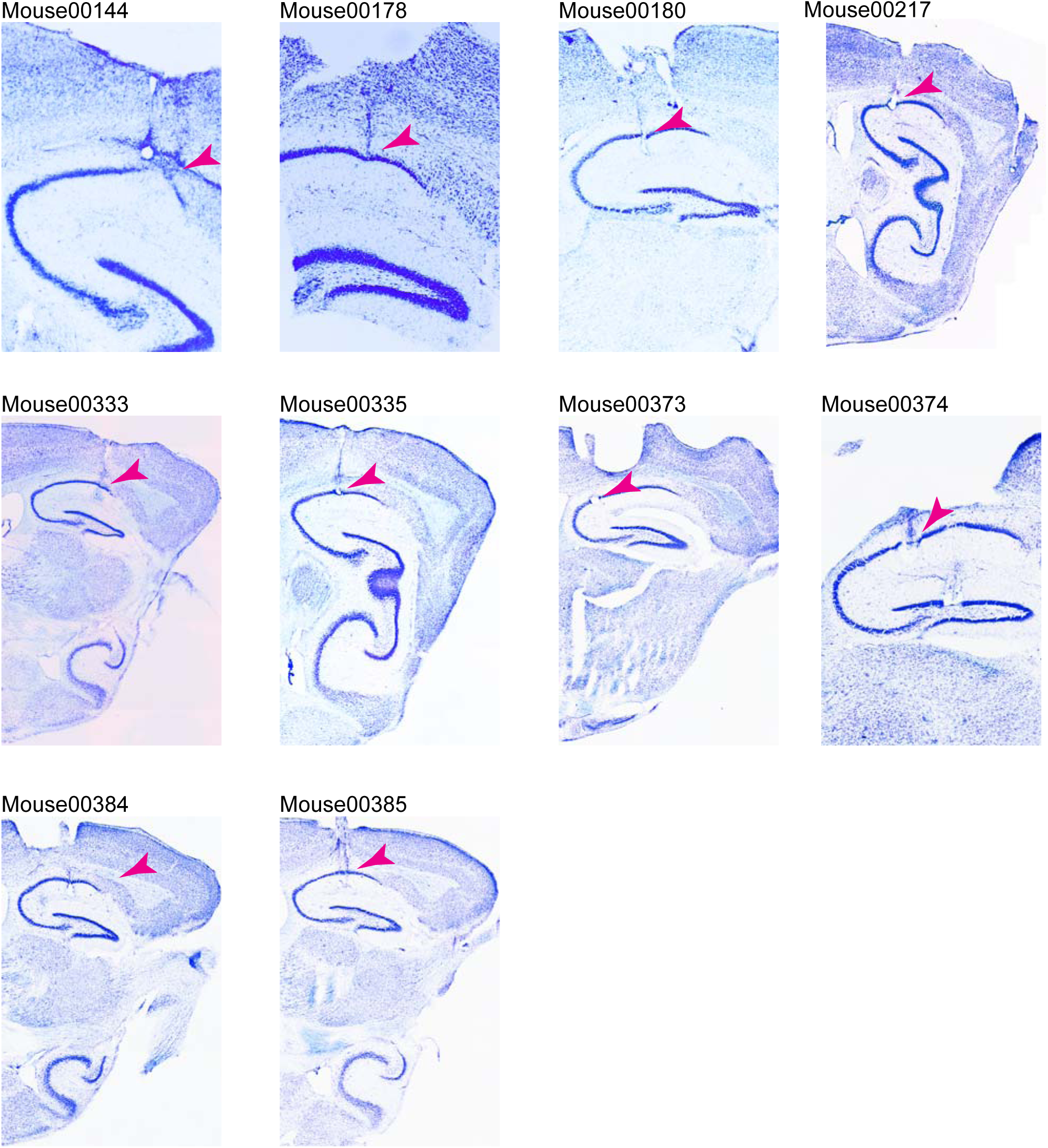
Histological validation of recording position in CA1 of WT and APP-KI mice. Brightfield images of cresyl violet stained sections of all animals recorded. Arrowhead points to the tip of tetrodes where the cells were recorded with electrolytic lesions made before sacrificing the animals.

**Fig. S4.**
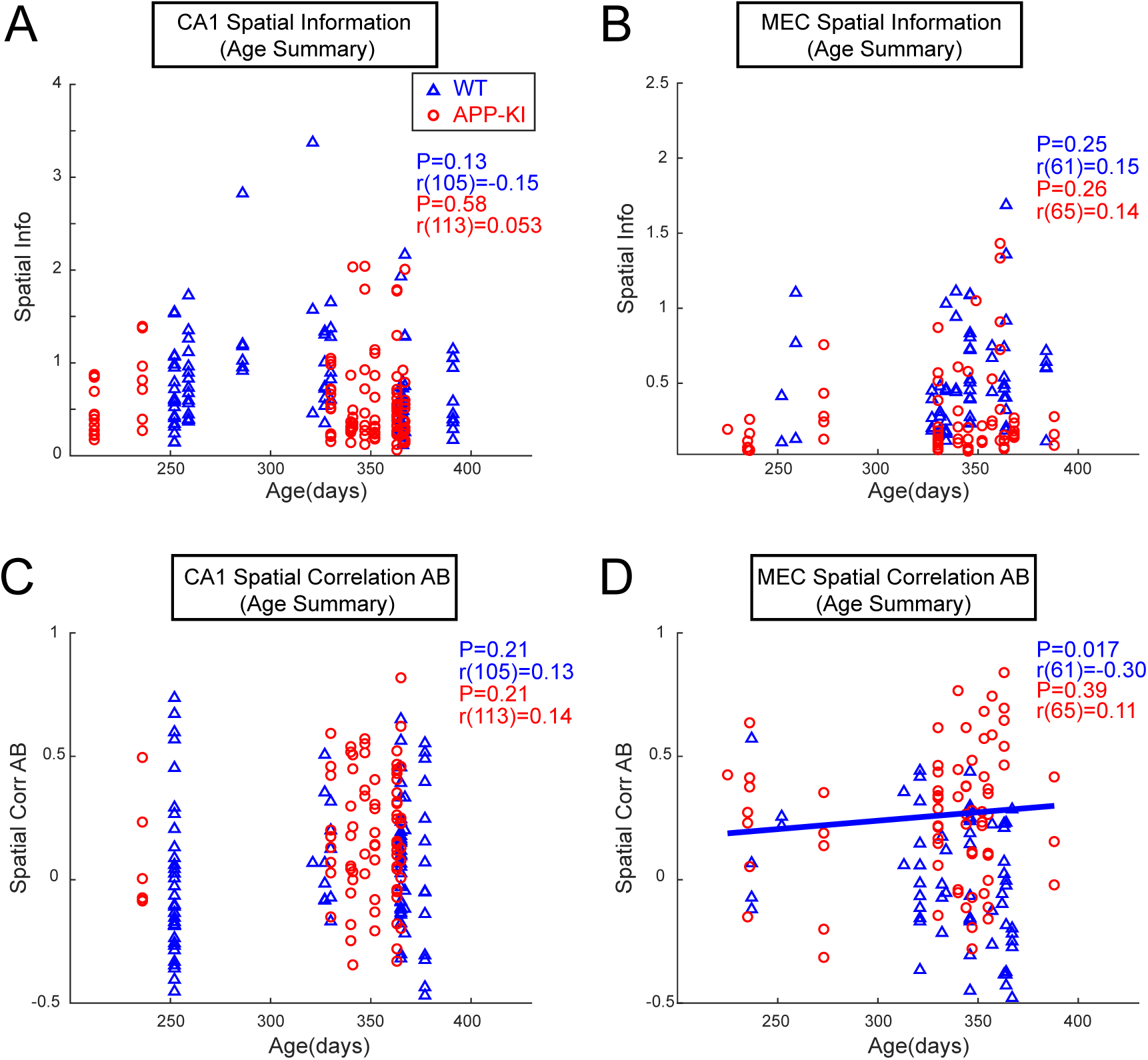
Spatial tuning and remapping property across age (7 - 13 months old) (**A-D**), Spatial information score and spatial correlation between Track A and Track B were plotted for CA1 neurons and MEC neurons across age (7-13 months old). No significant effects of aging were observed in APP-KI mice (p>0.05, Pearson correlation). A weak positive correlation was observed for spatial correlation in WT MEC neurons (r(61) = −0.30, p<0.05, Pearson correlation).

**Fig. S5.**
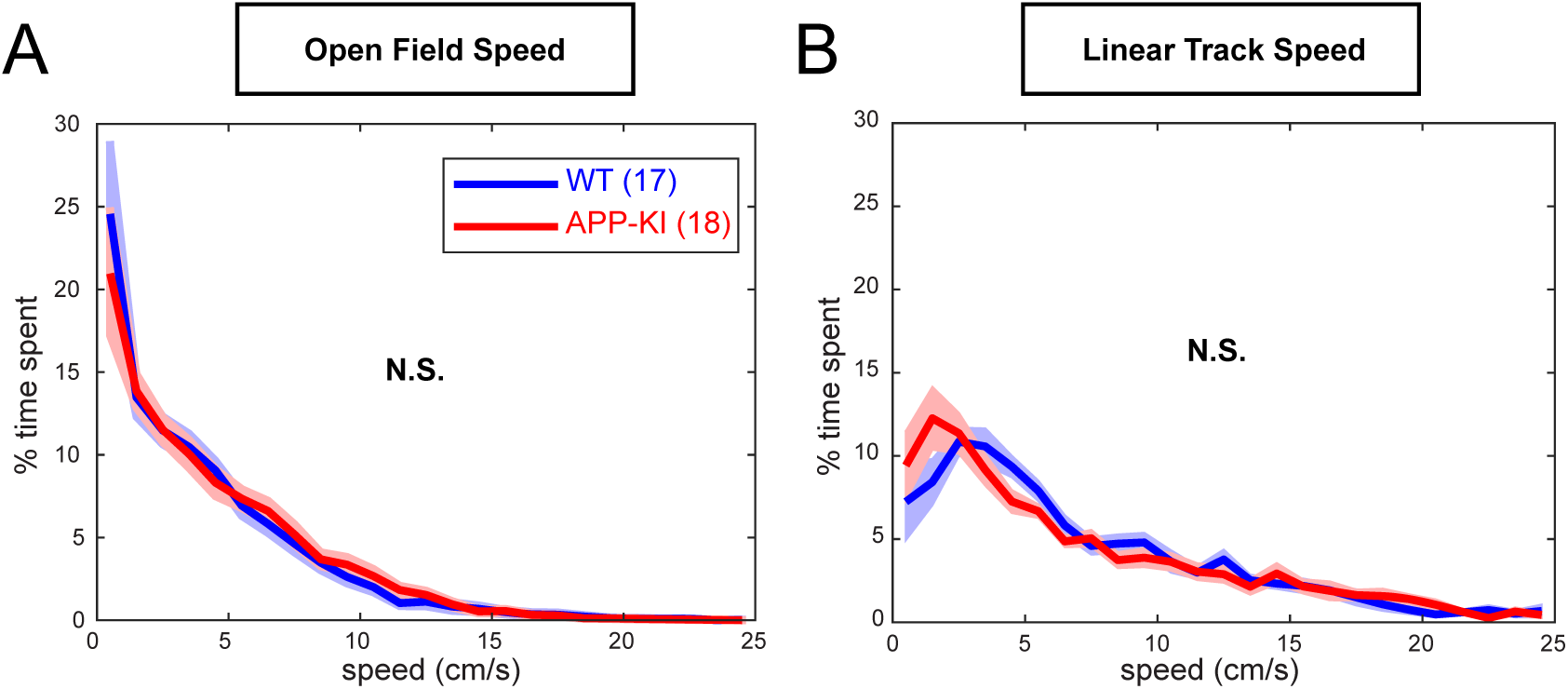
Running behaviors did not differ between WT and APP-KI mice. (**A**) Speed analysis during open field exploration. No significant difference between WT and APP-KI mice (q>0.05, FDR corrected for multiple comparison). (**B**) Speed analysis during linear track experiment. No significant difference between WT and APP-KI mice (q>0.05, FDR corrected for multiple comparison).

**Fig. S6.**
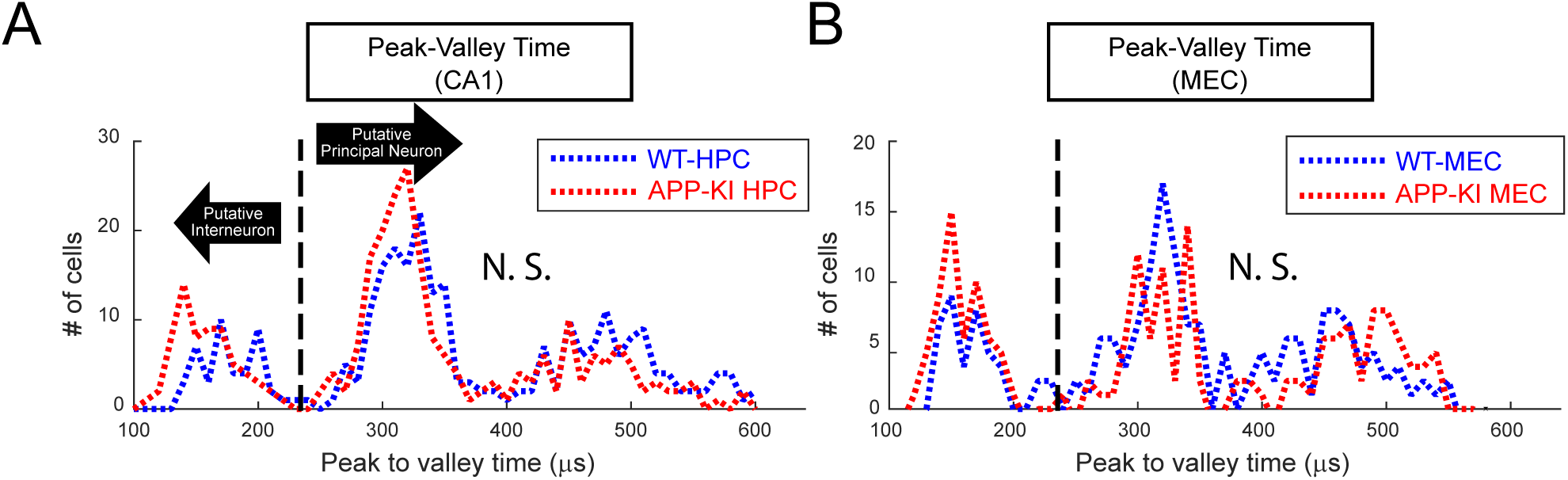
Electrophysiological features for classifying putative principal neurons in CA1 and MEC. (**A-B**) Peak to valley time of spike waveform was used to distinguish putative interneurons from principal neurons. Dashed line (230µs) represents cut off for interneuron to principal neuron classification (Bartho et al., 2004). (**A**) Peak to valley time histogram for neurons recorded from CA1. No difference was observed for the distribution of peak to valley time between WT and APP-KI mice (KS test, p >0.05). (**B**) Peak to valley time histogram for neurons recorded from MECs. No difference was observed for the distribution of peak to valley time between WT and APP-KI mice (KS test, p >0.05).

**Fig. S7.**
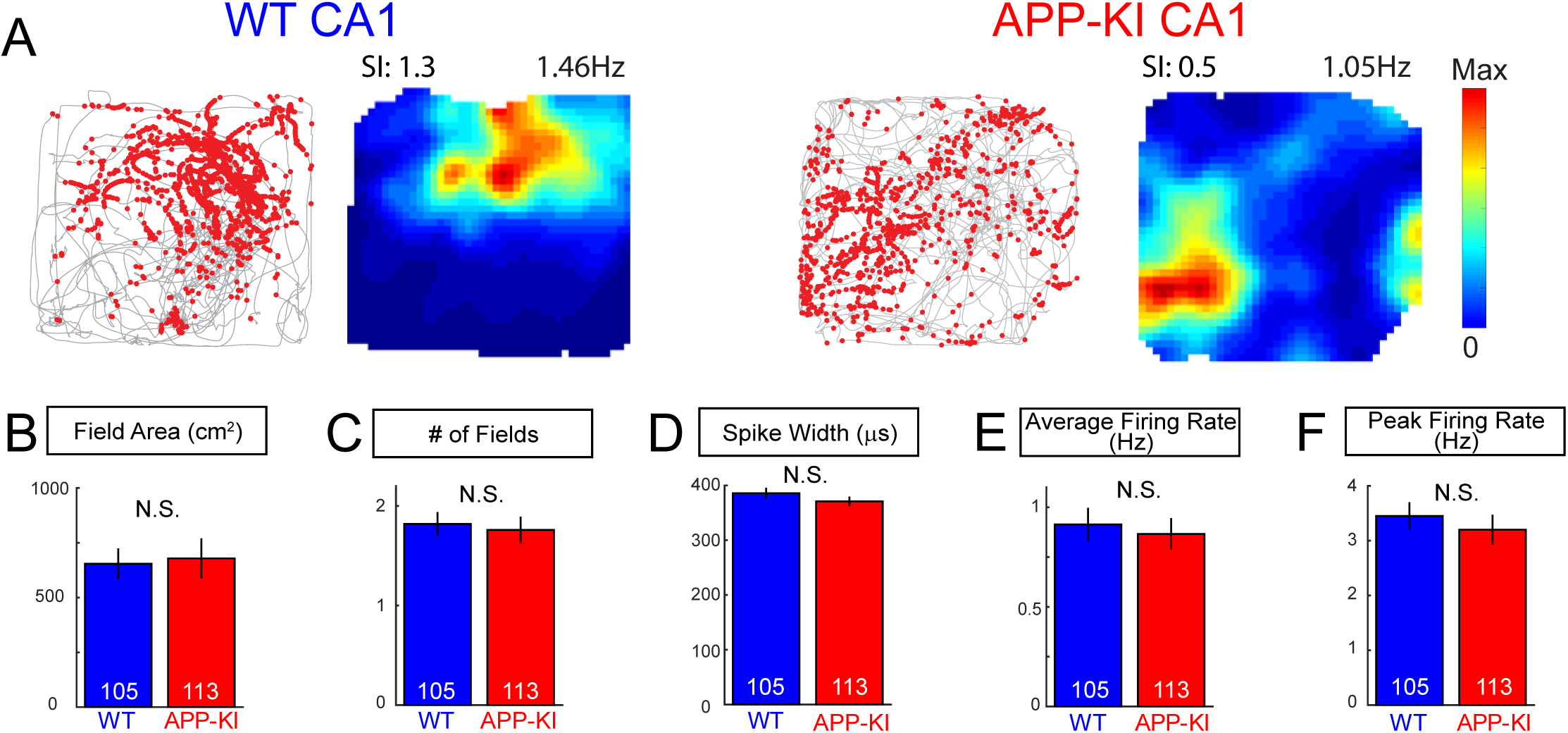
Additional properties of CA1 neurons during the open field task. (**A**) Representative CA1 neurons recorded from WT and APP-KI animals during open field task. Spatial information (SI) and average firing rate (Hz) for each cell is shown above rate map. (**B**) Field area (cm^2^) measured from CA1 neurons in open field task. (Wilcoxon rank sum test, p>0.05) (**C**) Number of fields measured from CA1 neurons in open field task. (Wilcoxon rank sum test, p>0.05) (**D**) Spike width measured from CA1 neurons in open field task. (Wilcoxon rank sum test, p>0.05) (**E**) Average firing rate measured from CA1 neurons in open field task. (Wilcoxon rank sum test, p>0.05) (**F**) Peak firing rate measured from CA1 neurons in open field task. (Wilcoxon rank sum test, p>0.05)

**Fig. S8.**
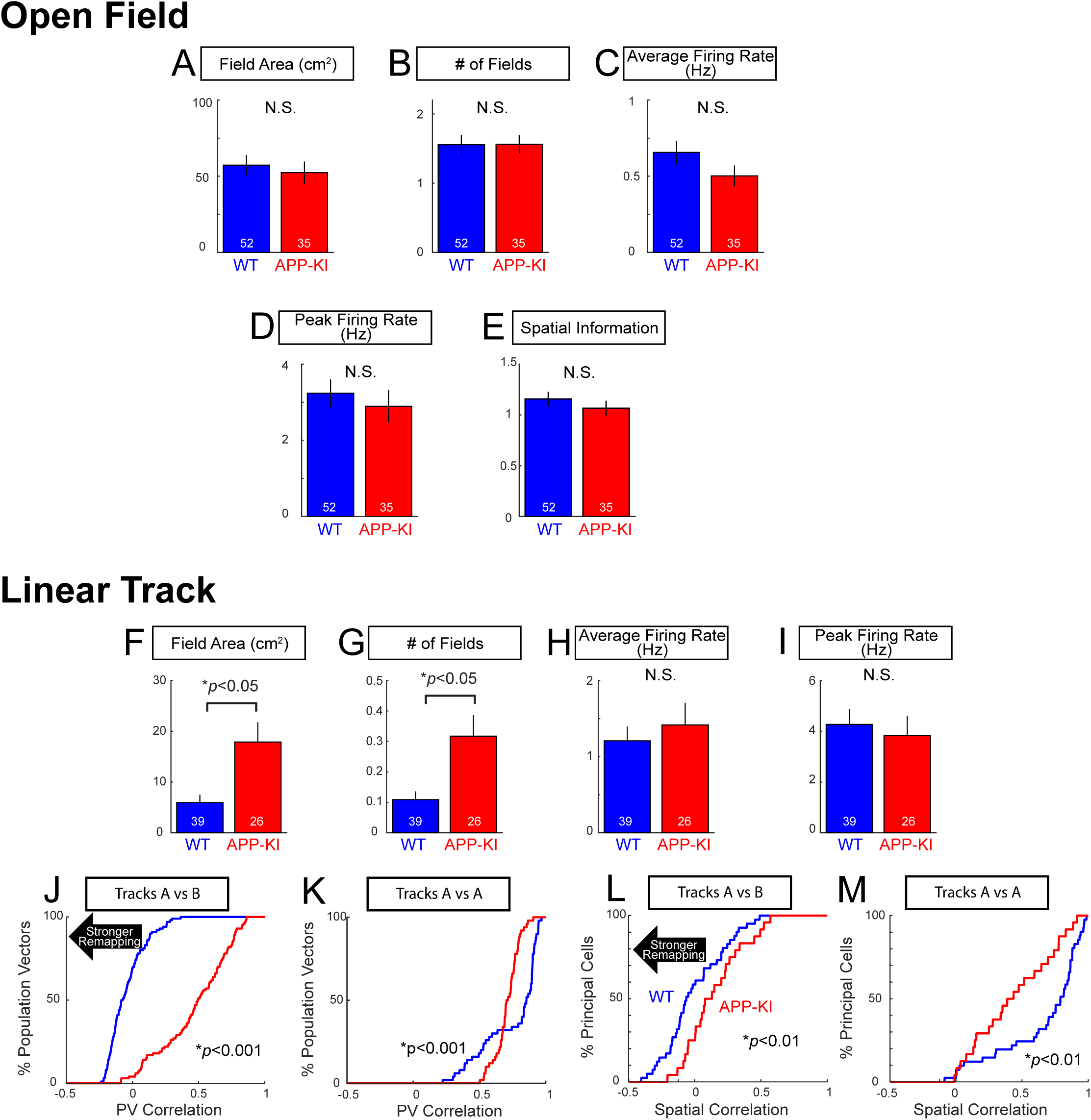
Properties of CA1 neurons defined as place cells in WT and APP-KI mice. (**A-E**) Properties of CA1 neurons defined as place cells in the open field task (n=52 cells in WT and n=35 cells in APP-KI mice). (**A**) Field area (cm^2^) measured from place cells of CA1 (Wilcoxon rank sum test, p>0.05). (**B**) Number of fields measured from place cells (Wilcoxon rank sum test, p>0.05). (**C**) Average firing rate measured from place cells (Wilcoxon rank sum test, p>0.05). (**D**) Peak firing rate measured from place cells (Wilcoxon rank sum test, p>0.05). (**E**) Spatial information measured from place cells. (Wilcoxon rank sum test, p>0.05). (**F-M**) Properties of CA1 neurons defined as place cells in the linear tracks (n=39 cells in WT and n=26 cells in APP-KI mice). (**F**) Field area (cm^2^) measured from place cells (Wilcoxon rank sum test, p<0.05). (**G**) Number of fields measured from place cells (Wilcoxon rank sum test, p<0.05). (**H**) Average firing rate measured from place cells (Wilcoxon rank sum test, p>0.05). (**I**) Peak firing rate measured from place cells (Wilcoxon rank sum test, p>0.05) (**J**) Cumulative distribution plot of PV correlation between Track A and Track B from place cells. Significant increase in PV correlation was observed compared to that of place cells in APP-KI mice (KS test, p <0.001). (**K**) Cumulative distribution plot of PV correlation between two recordings in Track A (KS test, p <0.001). (**L**) Cumulative distribution plot of Spatial Correlation between Track A and Track B (KS test, p<0.01). (**M**) Cumulative distribution plot of Spatial Correlation between two recordings in Track A (KS test, p <0.01).

**Fig. S9.**
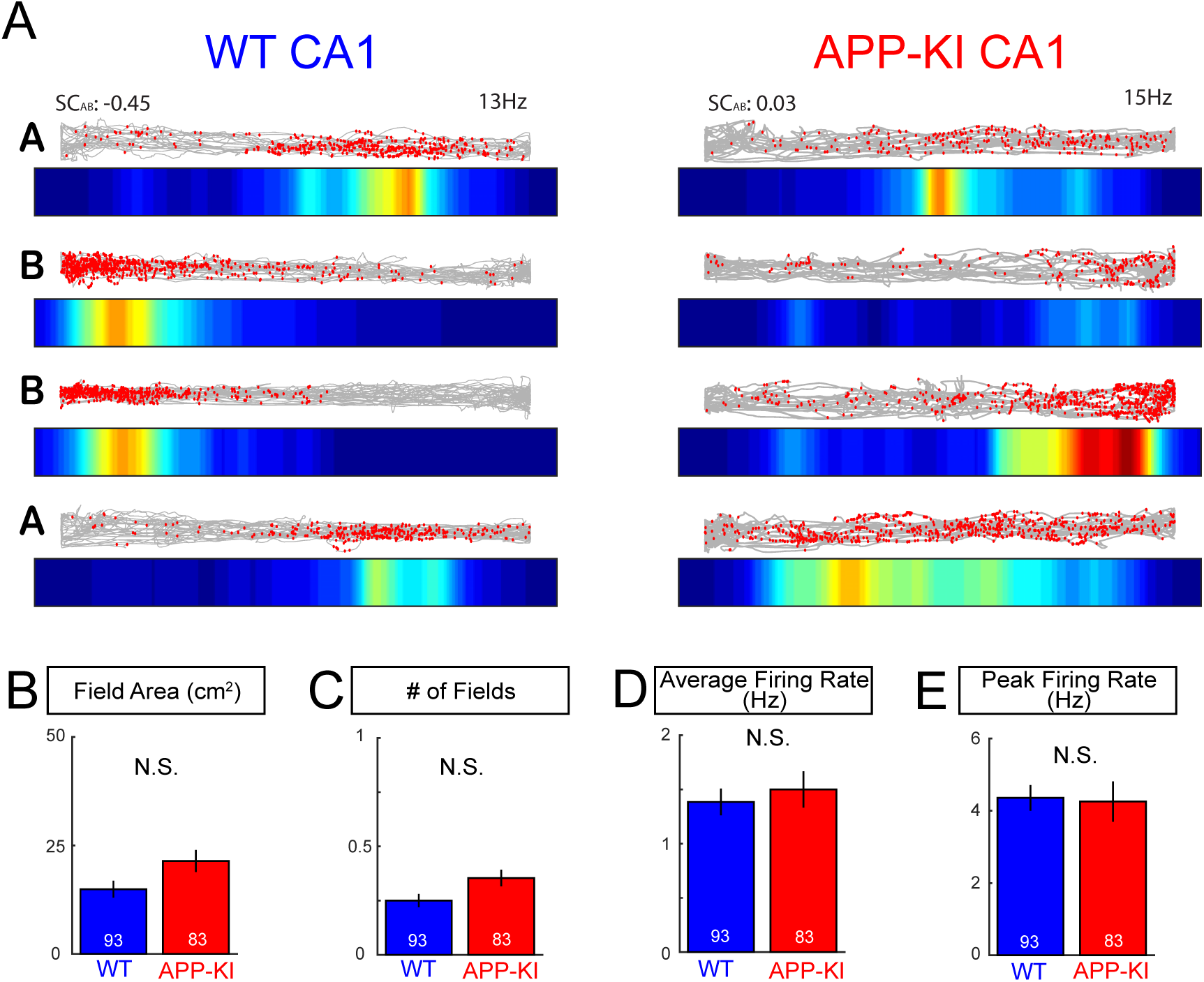
Additional properties of CA1 neurons during the remapping task. (**A**) Representative CA1 neurons recorded from WT and APP-KI mice during remapping task. Spatial correlation between Track A and Track B (SCAB) and average firing rate for each cell is shown above for each cell. (**B-G**) Properties of CA1 neurons in the linear tracks (**B**) Field area (cm^2^) (Wilcoxon rank sum, p>0.05). (**C**) Number of fields (Wilcoxon rank sum, p>0.05). (**D**) Average firing rate (Wilcoxon rank sum, p>0.05). (**E**) Peak firing rate (Wilcoxon rank sum, p>0.05).

**Fig. S10.**
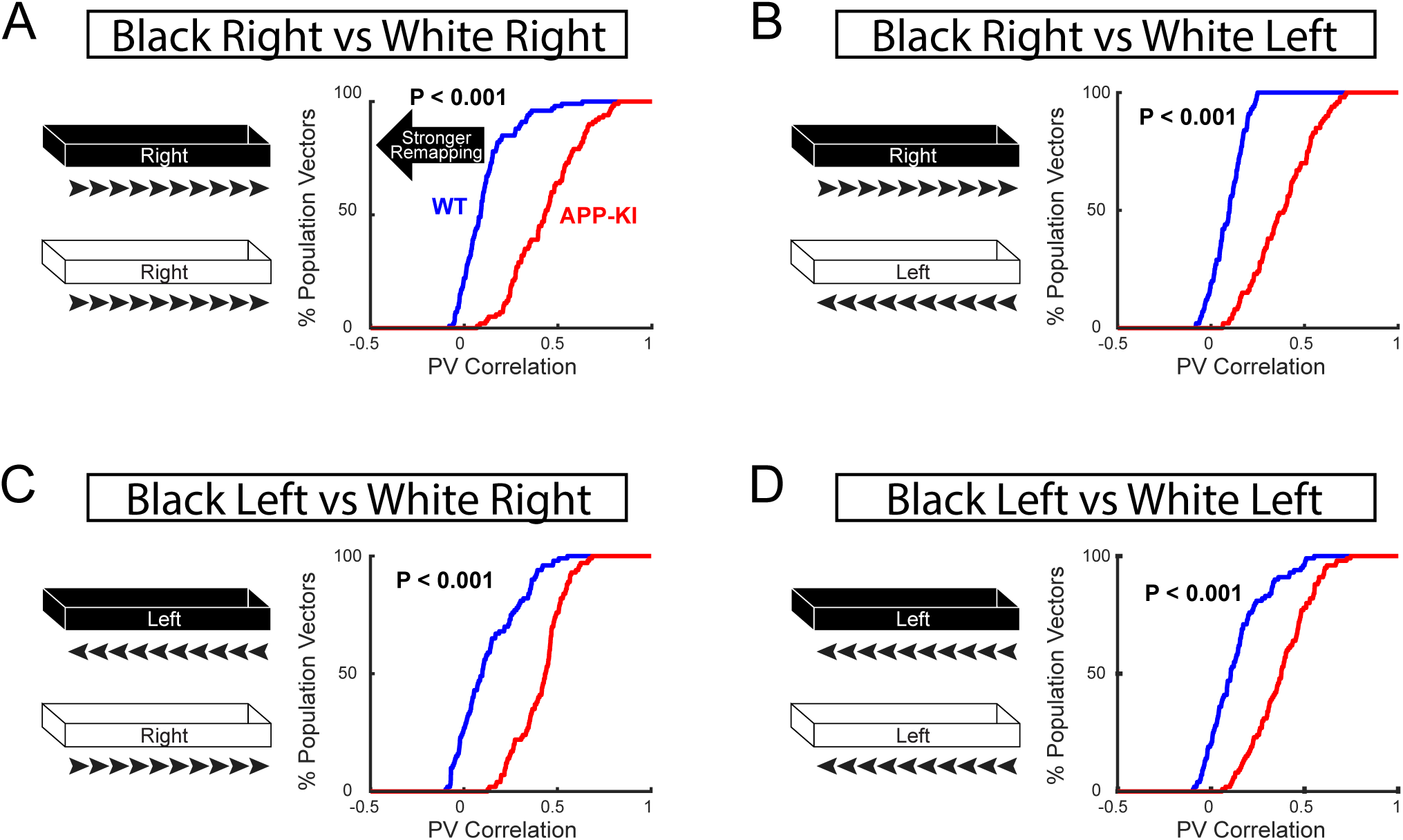
Disrupted remapping of CA1 neurons was observed regardless of animals’ running directions. (**A-D**) Cumulative distribution plot of PV Correlation between Track A and Track B from CA1 neurons during remapping task. Four possible combinations of directions have been analyzed. (**A**) Track A right vs Track B right. (**B**) Track A right vs Track B left. (**C**) Track A left vs Track B right. (**D**) Track A left vs Track B left. All combinations showed significant increase in PV correlation from CA1 neurons of APP-KI mice compared to that of WT mice indicating significant impairment in remapping regardless of directions (KS test, p <0.001 for **A-D**).

**Fig. S11.**
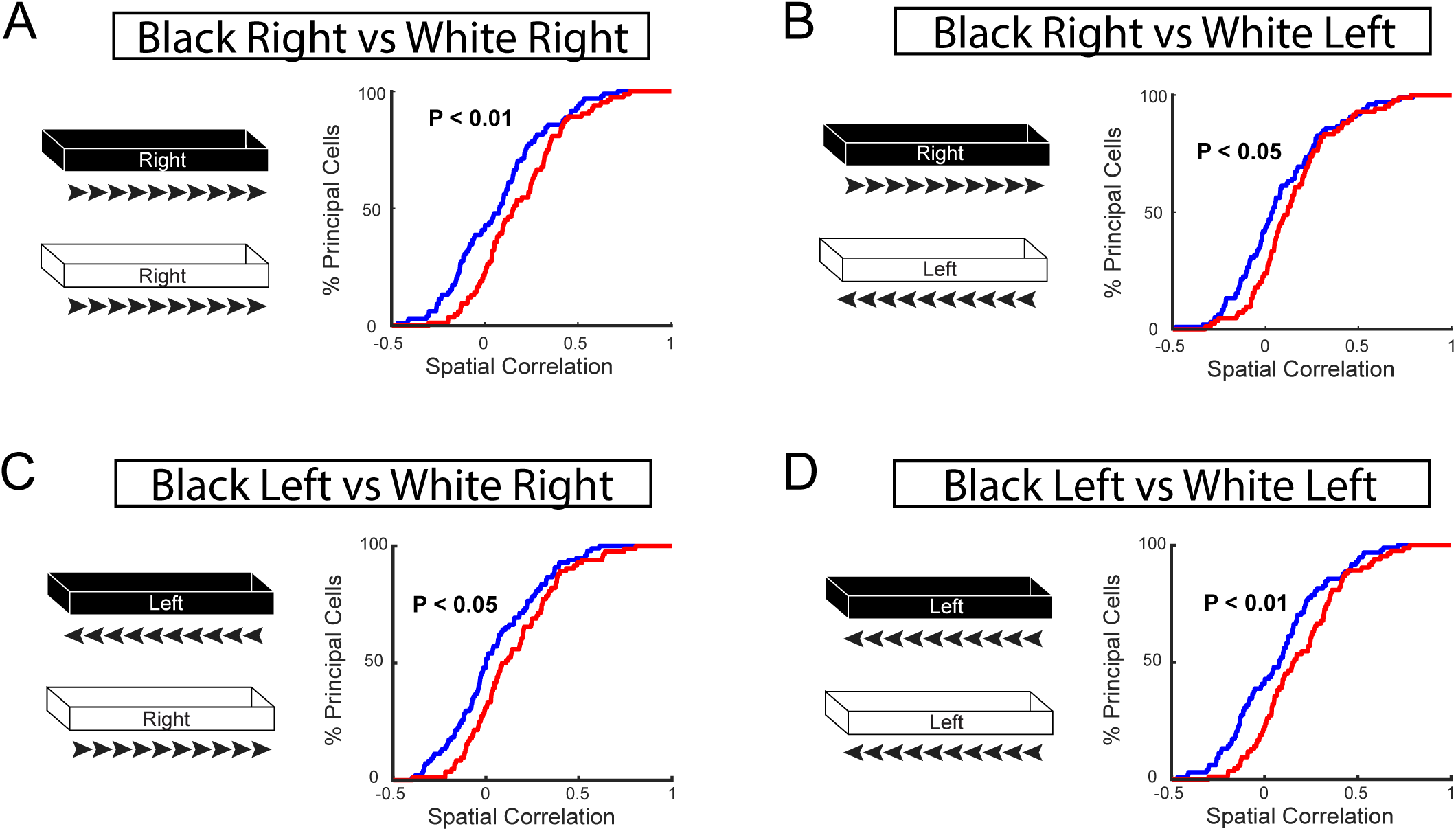
Disrupted spatial correlation of CA1 neurons was observed regardless of animals’ running directions. (**A-D**) Cumulative distribution plot of Spatial Correlation between Track A and Track B during remapping task. Four possible combinations of directions have been analyzed. (**A**) Track A right vs Track B right. (**B**) Track A right vs Track B left. (**C**) Track A left vs Track B right. (**D**) Track A left vs Track B left. All combinations showed significant increase in spatial correlation between Track A and Track B in CA1 neurons of APP-KI mice compared to that of WT mice, indicating significant impairment in remapping regardless of directions (KS test, p <0.05 or better for **A-D**).

**Fig. S12.**
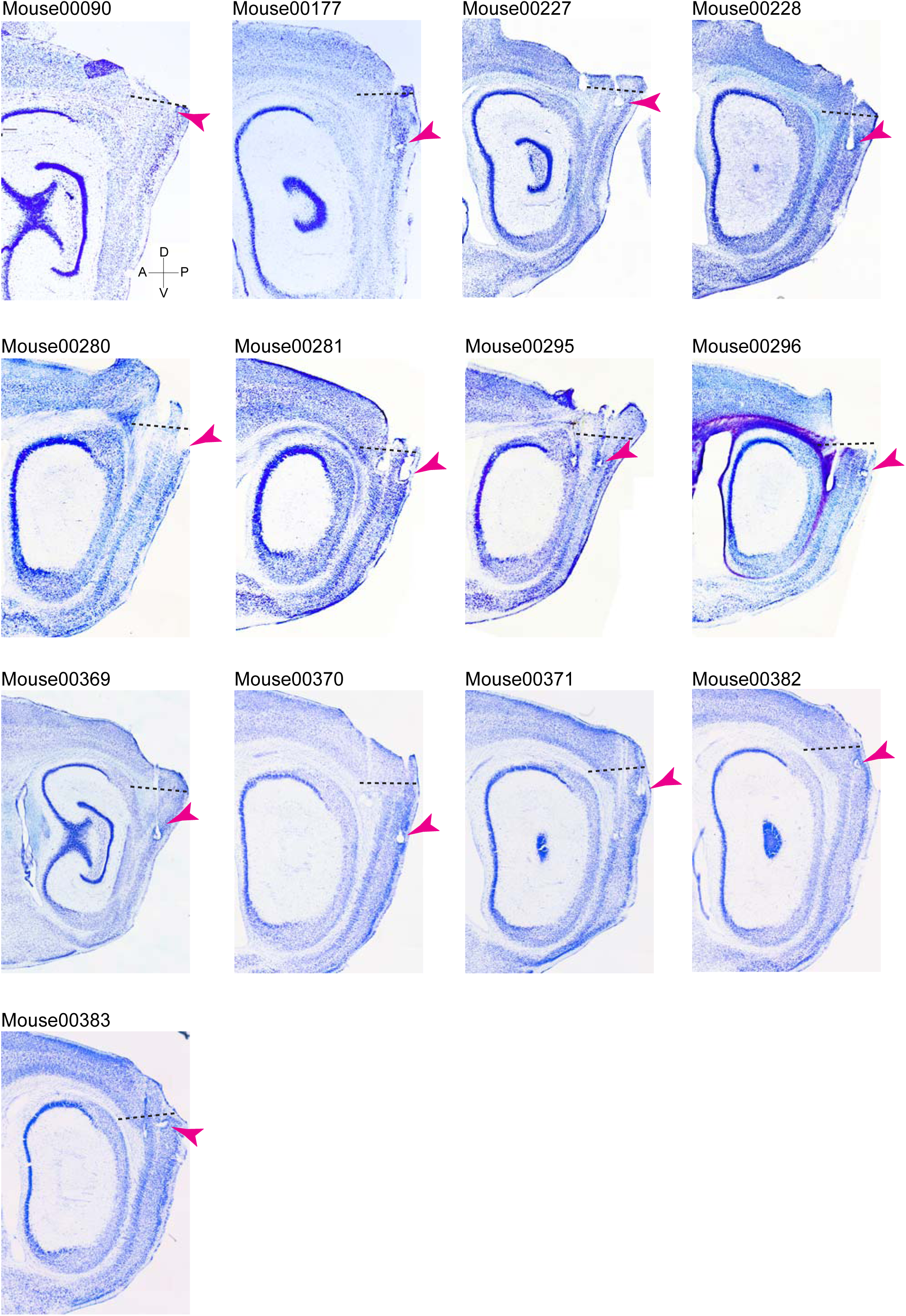
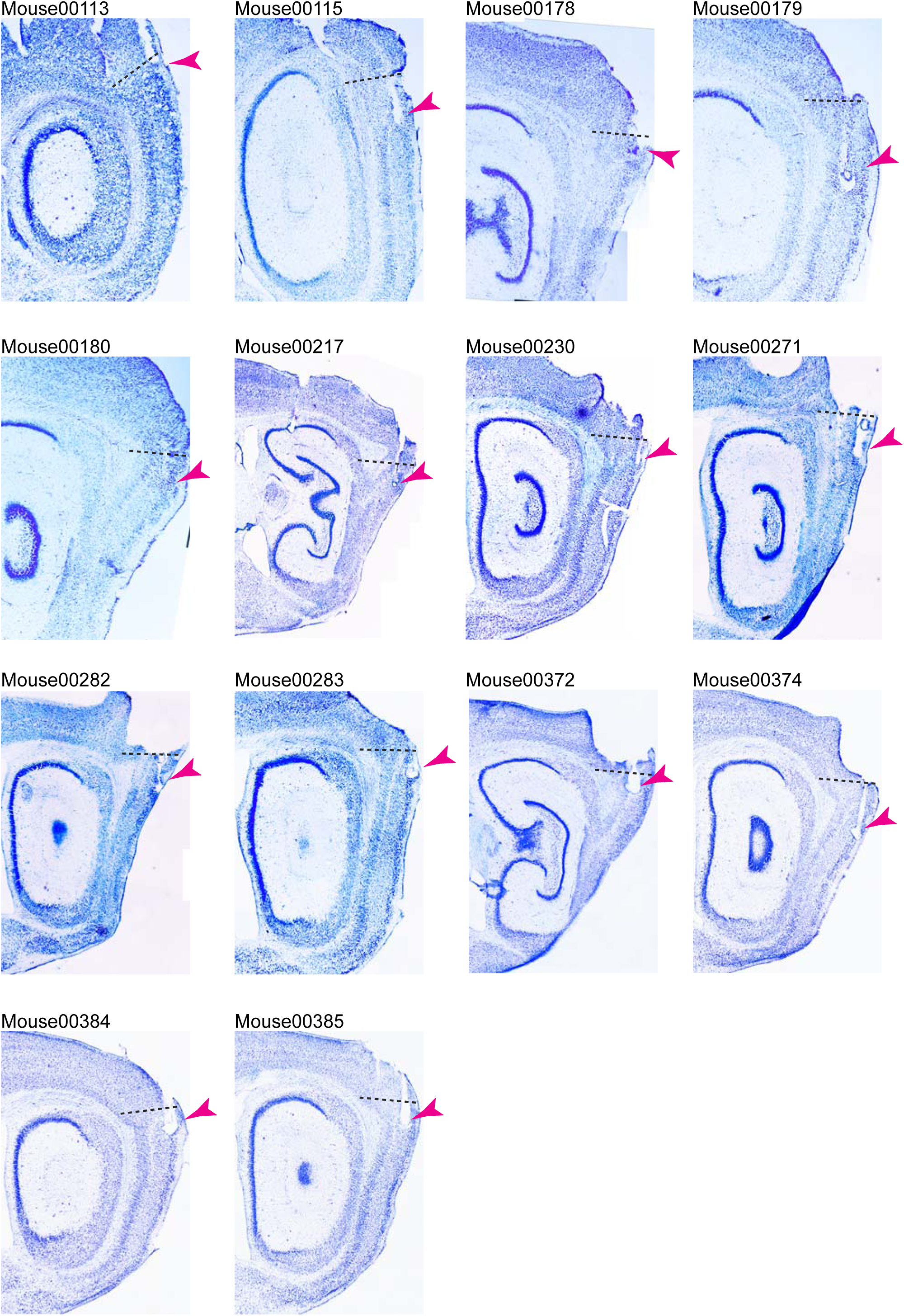
Histological validation of recording position in MEC of WT and APP-KI mice. Brightfield images of cresyl violet stained sections of all animals recorded. Arrowhead points to the tip of tetrodes where the cells were recorded with electrolytic lesions made before sacrificing the animals. Dashed line indicates the border between the MEC and the postrhinal cortex.

**Fig. S13.**
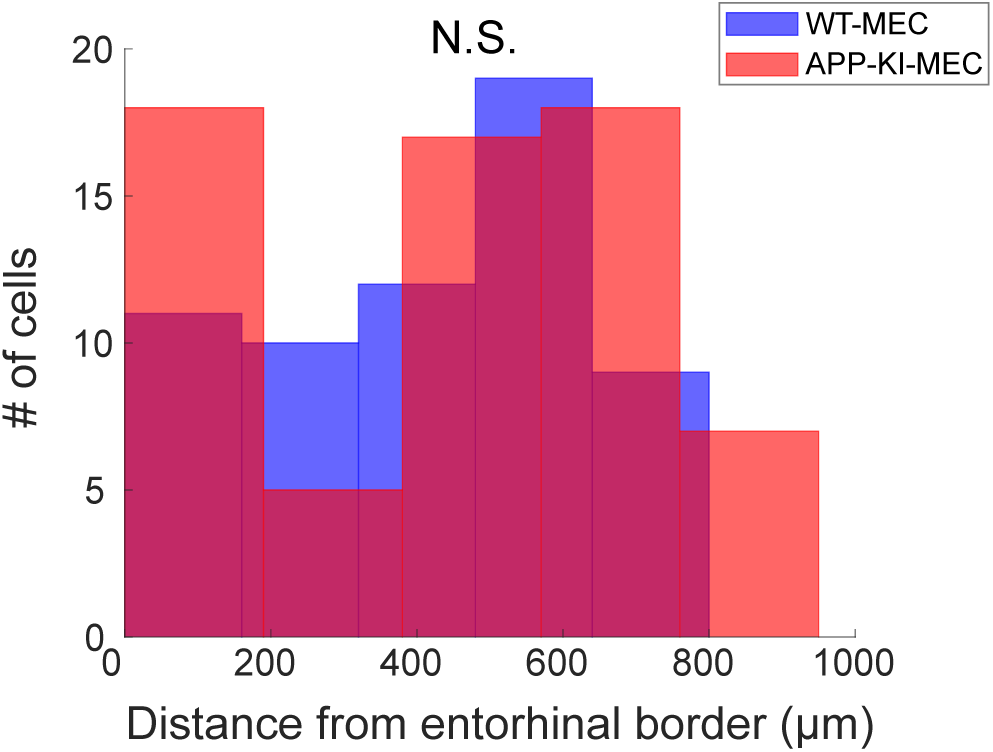
Recording positions in the MEC did not differ between WT and APP-KI mice. Recorded positions along dorsoventral axis of MEC. Distances were calculated from entorhinal border. The distribution of recording position along the dorsoventral axis did not differ between WT and APP-KI mice. (KS test, p>0.05).

**Fig. S14.**
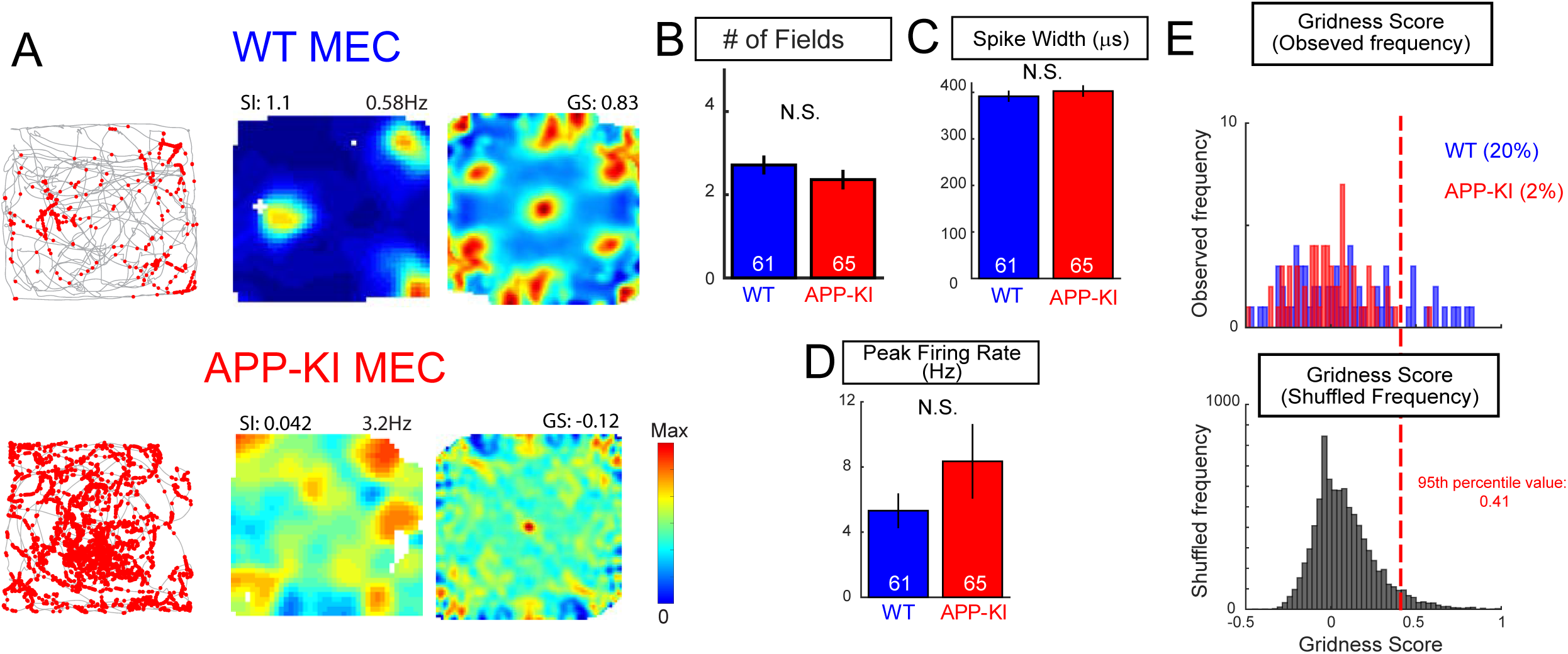
Additional properties of MEC neurons during the open field task. (**A**) Two representative MEC neurons recorded from WT mouse and APP-KI mouse during open field task. Spatial information (SI) and average firing rate (Hz) for each cell is shown above rate map. Gridness score (GS) for each cell is shown above autocorrelation map. (**B-D**) Properties of MEC neurons in open field task: (**B**) Number of fields (Wilcoxon rank sum test, p>0.05). (**C**) Spike width (Wilcoxon rank sum, p>0.05). (**D**) Peak firing rate (Wilcoxon rank sum test, p>0.05). (**E**) Top, distribution of gridness scores calculated from firing rate distributions of MEC neurons in the open field task (real data). Bottom, distribution of shuffled data based on 100 permutations per cell (shuffled data). Red dashed line indicates 95th percentile value (chance level) for a distribution based on all permutations. Percentage of cells that exceeded chance level is shown on top. 20% of MEC neurons in WT mice and 2% of MEC neurons in APP-KI mice passed this chance level, and they were considered as grid cells.

**Fig. S15.**
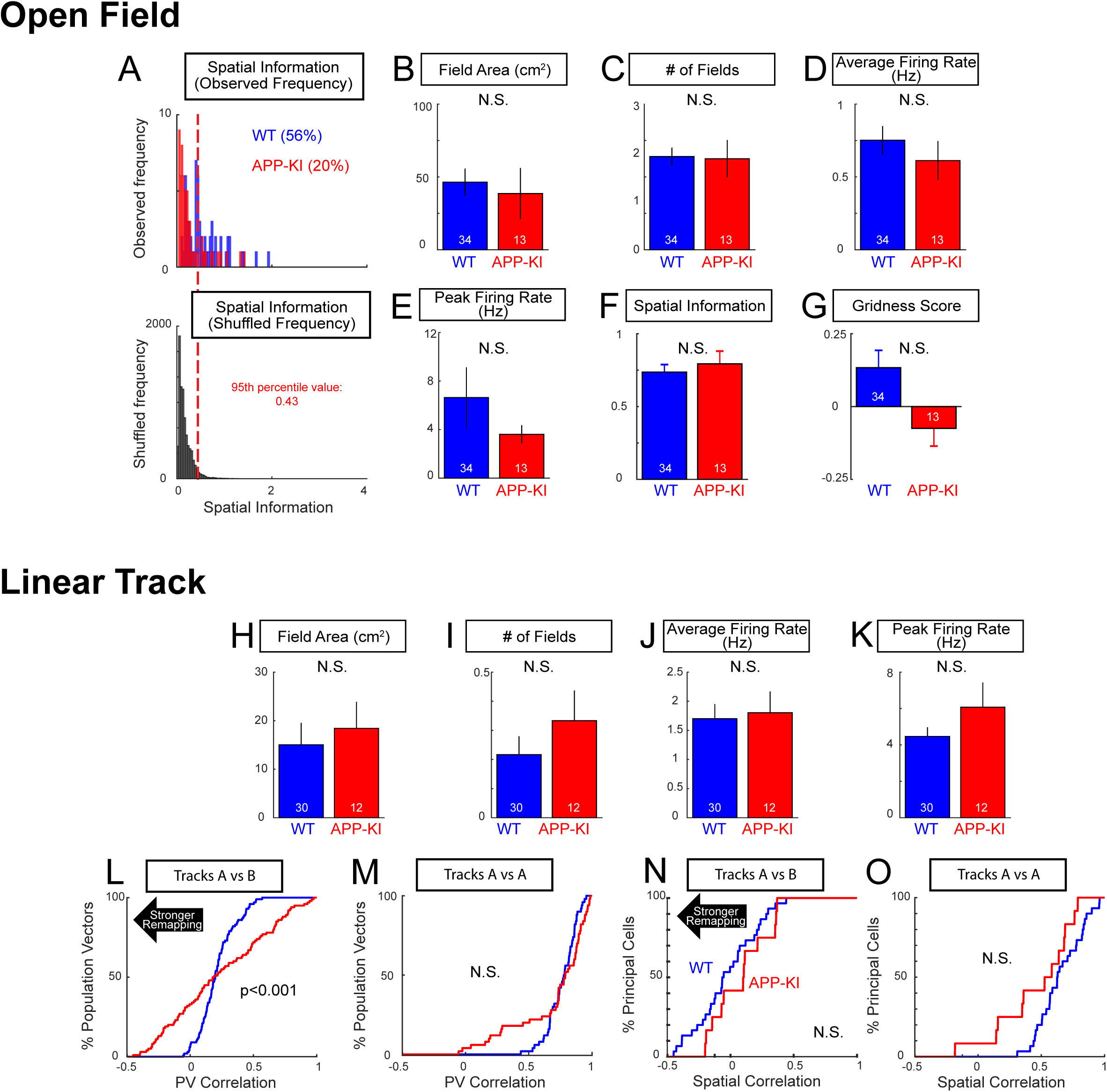
Properties of MEC neurons defined as spatially tuned cells. (**A**) Top, distribution of spatial information scores in all recorded MEC neurons in the open field task (real data). Bottom, distribution of shuffled data based on 100 permutations per cell (shuffled data). Red dashed line indicates 95th percentile value (chance level) for a distribution based on all permutations. Percentage of cells that exceeded chance level is shown on top. 56% of MEC neurons in WT mice and 20% of MEC neurons in APP-KI mice passed this chance level, and they were considered as “spatially tuned cells.” (**B-G**) Properties of MEC spatially tuned cells in the open field task. (**B**) Field area (cm^2^) (Wilcoxon rank sum test, p>0.05). (**C**) Number of fields (Wilcoxon rank sum test, p>0.05). (**D**) Average firing rate (Wilcoxon rank sum test, p>0.05). (**E**) Peak firing rate (Wilcoxon rank sum test, p>0.05). (**F**) Spatial information score (Wilcoxon rank sum test, p>0.05). (**G**) Gridness score (Wilcoxon rank sum test, p>0.05). (**H-O**) Properties of MEC spatially tuned cells in the linear tracks. (**H**) Field area (cm^2^) (Wilcoxon-rank-sum, p>0.05). (**I**) Number of fields (Wilcoxon rank sum test, p>0.05). (**J**) Average firing rate (Wilcoxon rank sum test, p>0.05). (**K**) Peak firing rate (Wilcoxon rank sum test, p>0.05). (**L**) Cumulative distribution plot of PV correlation between Track A and Track B (KS test, p<0.001). (**M**) Cumulative distribution plot of PV correlation between two recordings in Track A (KS test, p >0.05). (**N**) Cumulative distribution plot of spatial correlation between Track A and Track B (KS test, p >0.05). (**O**) Cumulative distribution plot of between two recordings in Track A (KS test, p>0.05).

**Fig. S16.**
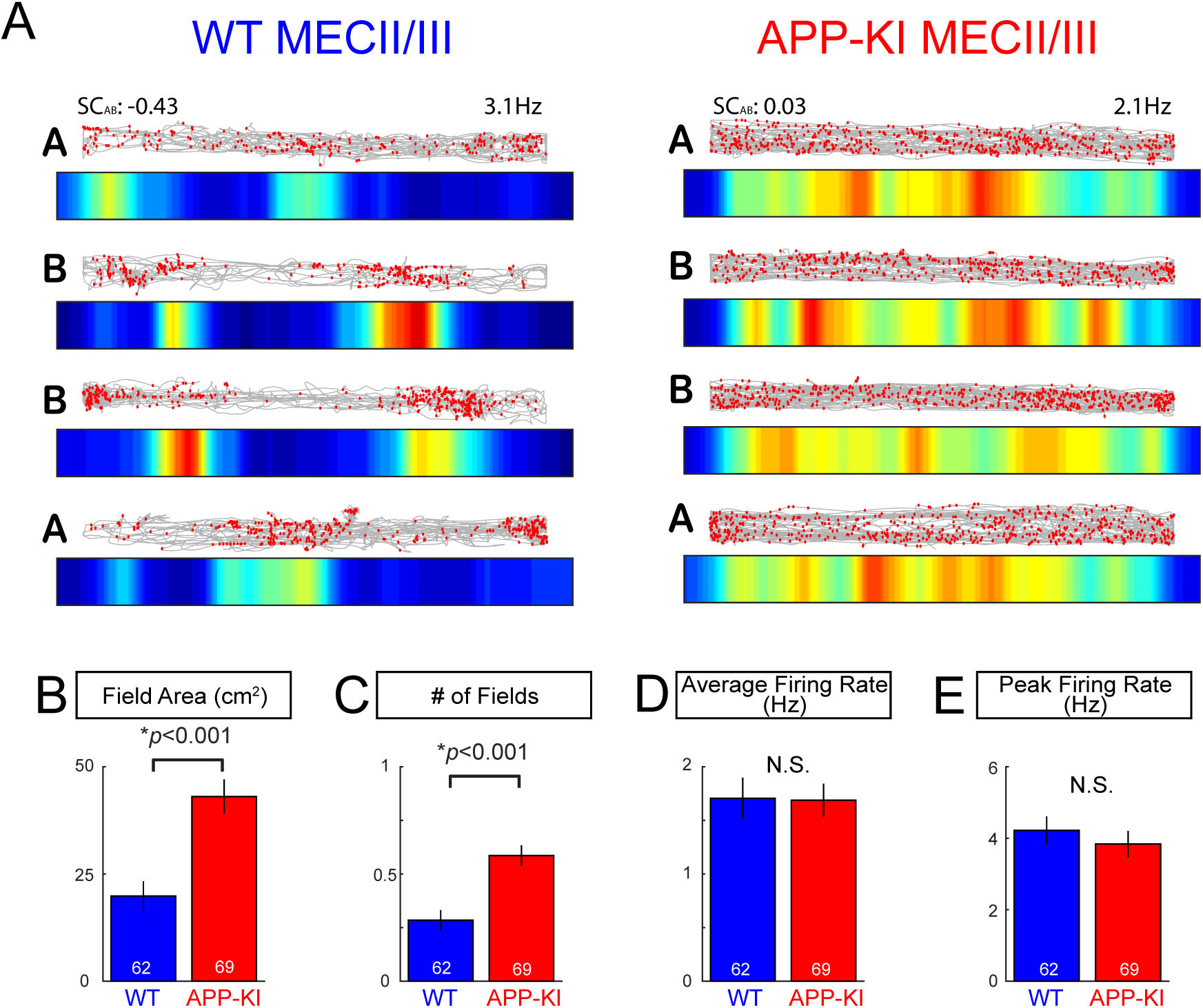
Additional properties of MEC neurons during the remapping task. (**A**) Two representative MEC neurons recorded from WT and APP-KI animals during the remapping task. Spatial correlation between Track A and Track B (SCAB) and average firing rate (Hz) for each cell is shown above for each cell. (**B**) Field area (cm^2^) (Wilcoxon-rank-sum, p<0.001). (**C**) Number of fields (Wilcoxon rank sum test, p<0.001). (**D**) Average firing rate (Wilcoxon-rank-sum, p>0.05). (**E**) Peak firing rate (Wilcoxon rank sum test, p>0.05).

**Fig. S17.**
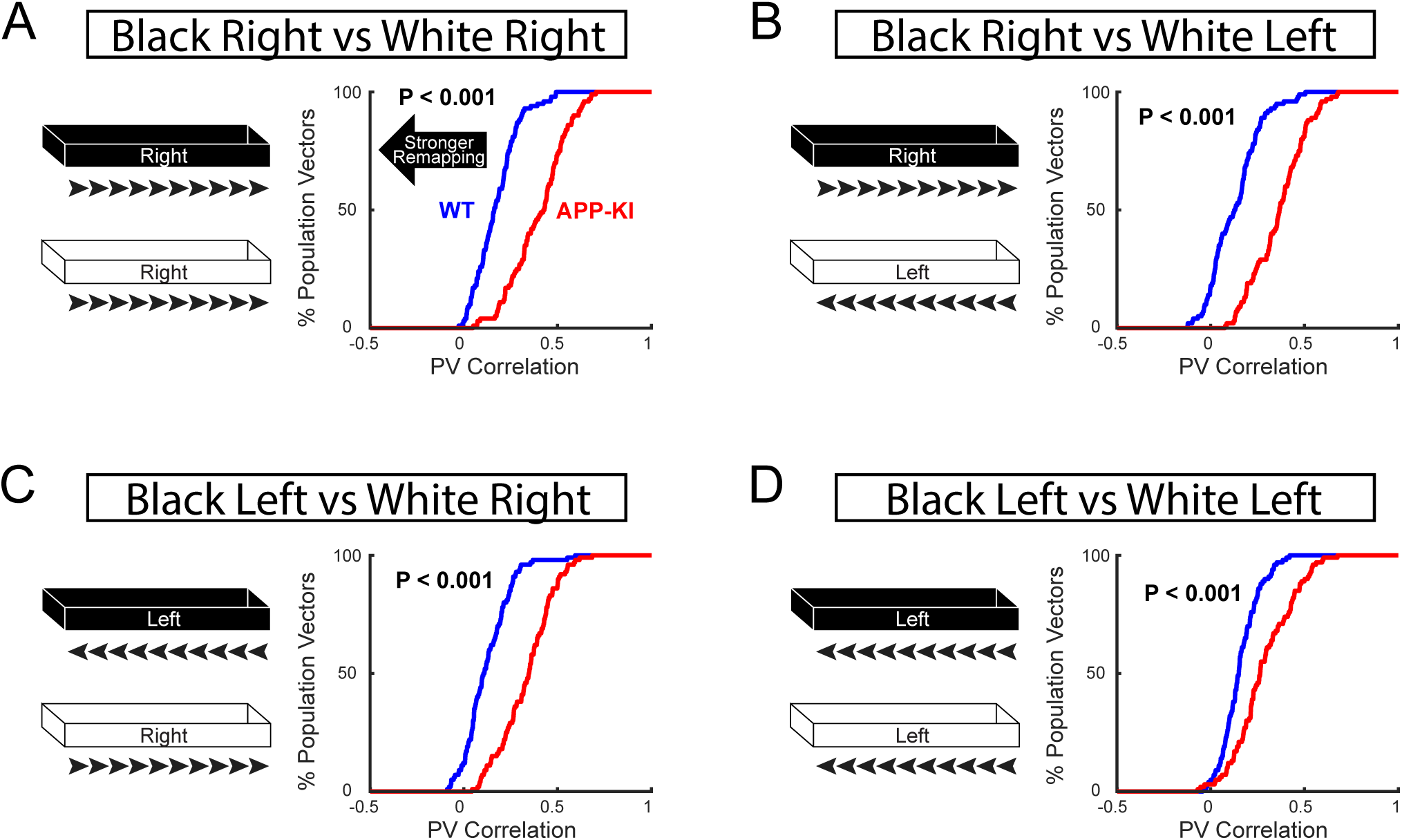
Disrupted remapping of MEC neurons was observed regardless of animals’ running directions. (**A-D**) Cumulative distribution plot of PV Correlation between Track A and Track B from MEC neurons during remapping task. Four possible combinations of directions have been analyzed. (**A**) Track A right vs Track B right. (**B**) Track A right vs Track B left. (**C**) Track A left vs Track B right. (**D**) Track A left vs Track B left. All combinations showed significant increase in PV correlation from MEC neurons of APP-KI mice compared to that of WT mice, indicating the impairment in remapping regardless of directions (KS test, p <0.001 for **A-D**).

**Fig. S18.**
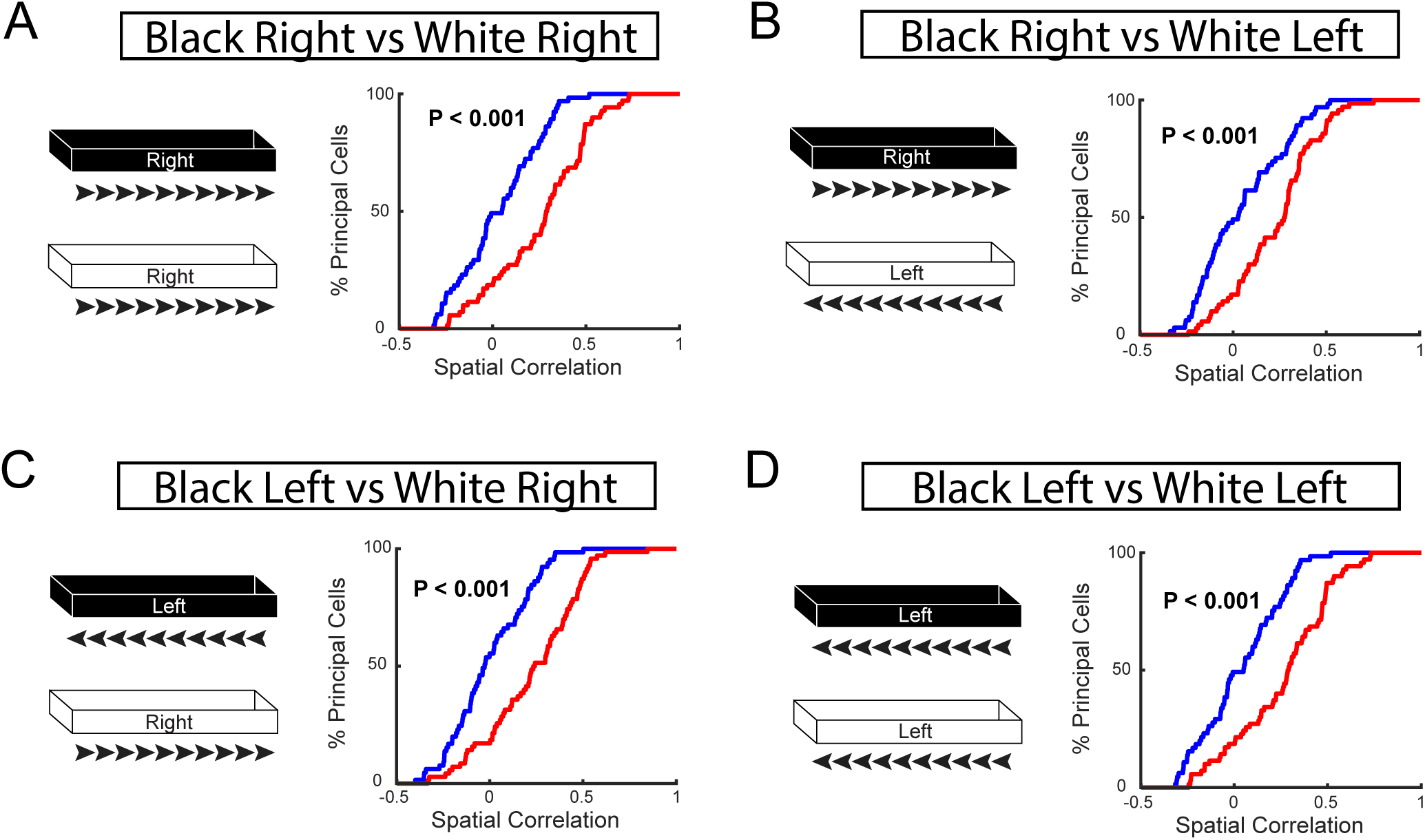
Disrupted spatial correlation of MEC neurons was observed regardless of animals’ running directions. (**A-D**) Cumulative distribution plot of spatial correlation AB (different tracks). Four possible combinations of directions have been analyzed. (**A**) Track A right vs Track B right. (**B**) Track A right vs Track B left. (**C**) Track A left vs Track B right. (**D**) Track A left vs Track B left. All combinations showed significant increase in Spatial Correlation AB from MEC neurons of APP-KI mice compared to that of WT mice, indicating significant impairment in remapping regardless of directions. (KS test, p <0.001 for **A-D**).

**Fig. S19.**
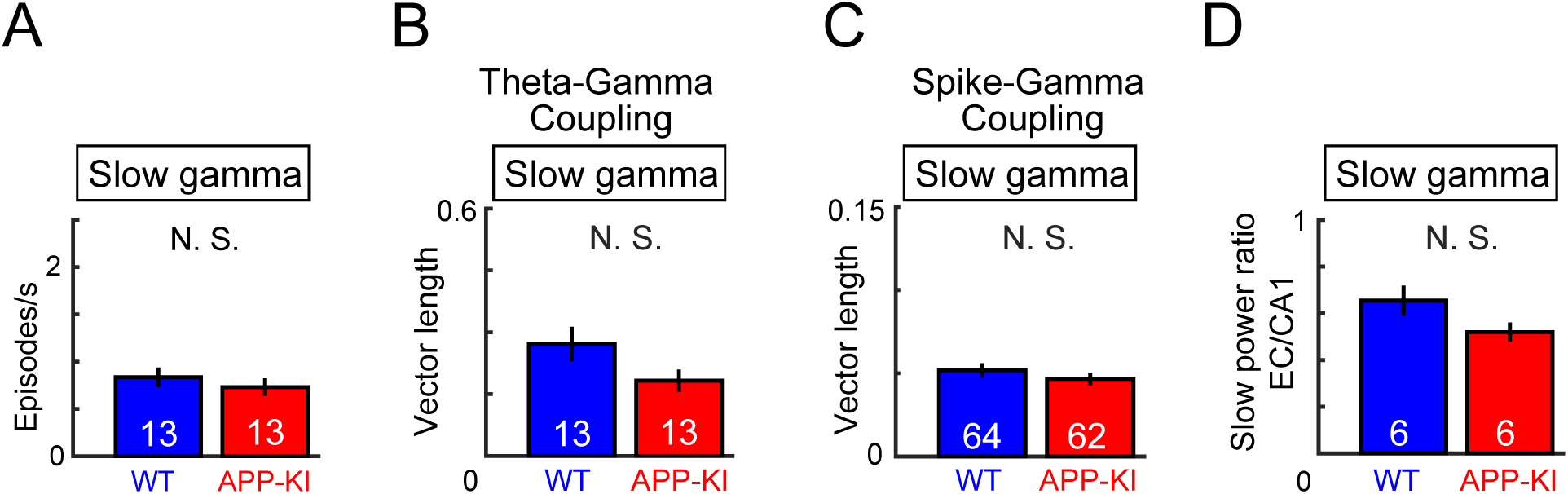
Slow gamma oscillations in the MEC. (**A**) Occurrence rates of slow gamma episodes in MEC for WT (n=13 mice) and APP-KI mice (n=13 mice) (Wilcoxon rank sum test, p >0.05). (**B**) Mean vector length for the distribution of slow gamma episodes across theta phase (Wilcoxon rank sum test, p>0.05). (**C**) Mean vector length for the distribution of spikes from WT MEC neurons (n=64 cells) and APP-KI MEC neurons (n=62 cells) across slow gamma phase (Wilcoxon ranks sum test, p>0.05). (**D**) MEC:CA1 ratio of slow gamma power during slow gamma episodes detected in CA1 (Wilcoxon rank sum test, p >0.05).

**Fig. S20.**
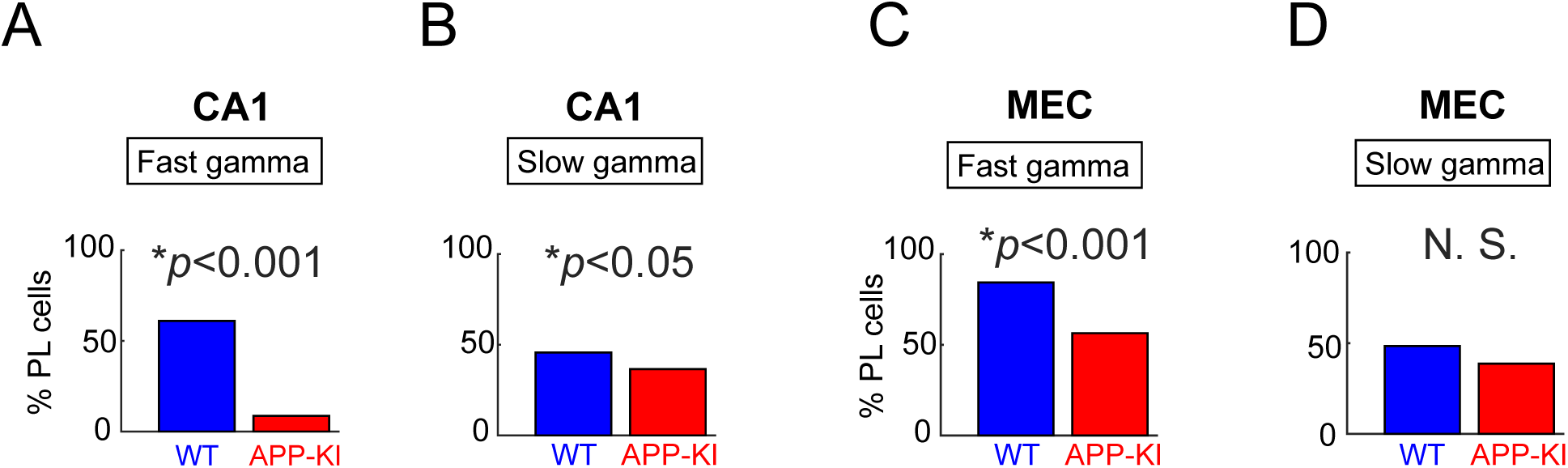
Percentage of neurons showing spike phase locking to slow and fast gamma oscillations. (**A**) CA1 neurons’ spikes phase locking to fast gamma oscillations (Binomial test, p<0.001) (**B**) CA1 neurons’ spikes phase locking to slow gamma oscillations (Binomial test, p<0.05) (**C**) MEC neurons’ spikes phase locking to fast gamma oscillations (Binomial test, p<0.001) (**D**) MEC neurons’ spikes phase locking to slow gamma oscillations (Binomial test, p>0.05)

**Table S1.** Distribution of recorded principal cells across all animals. The total number of isolated CA1 and MEC layer 2/3 cells recorded from each animal.

Distribution of recorded principal cells across all animals
| Animal ID | Genotype | Gender | Drive Used | Recorded Age | Recorded Area | Number of Principal Cells Recorded (open field) | Number of Principal Cells Recorded (linear track) |
| --- | --- | --- | --- | --- | --- | --- | --- |
| mouse00113 | APP-KI | Male | MEC only | 252 | MEC Layer 2/3 | 1 | 1 |
| mouse00115 | APP-KI | Male | MEC only | 237 | MEC Layer 2/3 | 5 | 5 |
| mouse00144 | APP-KI | Male | HPC only | 212 | CA1 | 12 | N/A |
| mouse00178 | APP-KI | Male | Split drive | 353 | CA1<br>MEC Layer 2/3 | 11<br>4 | 10<br>3 |
| mouse00179 | APP-KI | Male | Split drive | 236 | MEC Layer 2/3 | 3 | 3 |
| mouse00180 | APP-KI | Female | Split Drive | 236 | CA1<br>MEC Layer 2/3 | 7<br>4 | 7<br>4 |
| mouse00217 | APP-KI | Female | Split Drive | 377 | CA1<br>MEC Layer 2/3 | 26<br>5 | 24<br>5 |
| mouse00230 | APP-KI | Female | Split Drive | 365 | MEC Layer 2/3 | 3 | 3 |
| mouse00271 | APP-KI | Male | MEC only | 368 | MEC Layer 2/3 | 6 | 4 |
| mouse00282 | APP-KI | Female | MEC only | 368 | MEC Layer 2/3 | 6 | 8 |
| mouse00283 | APP-KI | Male | MEC only | 371 | MEC Layer 2/3 | 3 | 2 |
| mouse00333 | APP-KI | Male | HPC only | 380 | CA1 | 6 | 4 |
| mouse00335 | APP-KI | Male | HPC only | 379 | CA1 | 13 | 7 |
| mouse00372 | APP-KI | Male | Split drive | 358 | MEC Layer 2/3 | 1 | 5 |
| mouse00373 | APP-KI | Male | Split drive | 359 | CA1 | 10 | 9 |
| mouse00374 | APP-KI | Male | Split drive | 357 | CA1<br>MEC Layer 2/3 | 9<br>5 | 7<br>7 |
| mouse00384 | APP-KI | Male | Split drive | 371 | CA1<br>MEC Layer 2/3 | 6<br>7 | 6<br>6 |
| mouse00385 | APP-KI | Male | Split drive | 371 | CA1<br>MEC Layer 2/3 | 13<br>13 | 9<br>13 |

Distribution of recorded principal cells across all animals
| Animal ID | Genotype | Gender | Drive Used | Recorded Age | Recorded Area | Number of Principal Cells<br>Recorded (open field) | Number of Principal Cells<br>Recorded (linear track) |
| --- | --- | --- | --- | --- | --- | --- | --- |
| mouse00090 | WT | Male | Split Drive | 362 | MEC Layer 2/3 | 5 | 2 |
| mouse00106 | WT | Male | HPC only | 286 | CA1 | 6 | N/A |
| mouse00177 | WT | Male | Split Drive | 327 | CA1 | 8 | 6 |
|  |  |  |  |  | MEC Layer 2/3 | 4 | 7 |
| mouse00227 | WT | Female | Split Drive | 259 | CA1 | 15 | 13 |
|  |  |  |  |  | MEC Layer 2/3 | 3 | 4 |
| mouse00228 | WT | Female | Split Drive | 259 | CA1 | 22 | 21 |
|  |  |  |  |  | MEC Layer 2/3 | 2 | 2 |
| mouse00229 | WT | Female | Split Drive | 390 | CA1 | 11 | 14 |
| mouse00280 | WT | Male | MEC only | 371 | MEC Layer 2/3 | 3 | 3 |
| mouse00281 | WT | Male | MEC only | 369 | MEC Layer 2/3 | 5 | 5 |
| mouse00295 | WT | Male | MEC only | 370 | MEC Layer 2/3 | 7 | 7 |
| mouse00296 | WT | Female | MEC only | 353 | MEC Layer 2/3 | 14 | 14 |
| mouse00331 | WT | Male | HPC only | 374 | CA1 | 11 | 9 |
| mouse00332 | WT | Male | HPC only | 375 | CA1 | 18 | 19 |
| mouse00369 | WT | Male | Split Drive | 339 | CA1 | 3 | 1 |
|  |  |  |  |  | MEC Layer 2/3 | 4 | 4 |
| mouse00370 | WT | Female | Split Drive | 356 | CA1 | 5 | 5 |
|  |  |  |  |  | MEC Layer 2/3 | 5 | 6 |
| mouse00371 | WT | Male | Split Drive | 327 | MEC Layer 2/3 | 1 | 1 |
| mouse00382 | WT | Male | Split Drive | 322 | MEC Layer 2/3 | 6 | 6 |
| mouse00383 | WT | Female | Split Drive | 320 | CA1 | 6 | 5 |
|  |  |  |  |  | MEC Layer 2/3 | 2 | 2 |

**Table S2.**
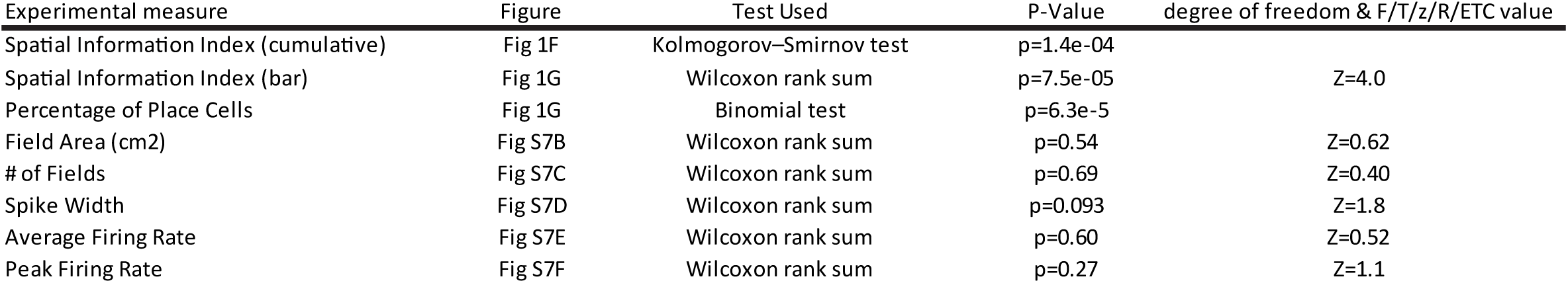
Statistics used for properties of CA1 neurons from open field. Summary of statistics applied on analysis from CA1 neurons recorded during open field task. (WT CA1 neurons=105 and APP-KI CA1 neurons=113)

**Table S3.** Statistics used for properties of CA1 neurons from linear track. Summary of statistics applied on analysis from CA1 neurons recorded during open field task. (WT CA1 neurons=93 and APP-KI CA1 neurons=83)

CA1 neurons from linear track
| WT CA1 neurons (n=93) vs APP-KI CA1 neurons (n=83) |  |  |  |  |
| --- | --- | --- | --- | --- |
| Experimental measure | Figure | Test Used | P-Value | degree of freedom & F/T/z/R/ETC value |
| Population Vector Correlation AB | Fig 1L | Wilcoxon rank sum | p=4.1e-29 | Z=11 |
| Population Vector Correlation AA | Fig 1L | Wilcoxon rank sum | p=0.42 | Z=0.81 |
| Field Area | Fig S9B | Wilcoxon rank sum | p=0.10 | Z=1.7 |
| # of Fields | Fig S9C | Wilcoxon rank sum | p=0.070 | Z=1.8 |
| Average Firing Rate | Fig S9D | Wilcoxon rank sum | p=0.52 | Z=0.65 |
| Peak Firing Rate | Fig S9E | Wilcoxon rank sum | p=0.10 | Z=1.6 |
| Population Vector Correlation AA | Fig S9F | Wilcoxon rank sum | p=0.42 | Z=0.81 |
| Population Vector Correlation AB | Fig S9G | Wilcoxon rank sum | p=4.1e-29 | Z=11 |
| Population Vector Correlation ArBr | Fig S10A | Kolmogorov–Smirnov test | p=1.8e-26 |  |
| Population Vector Correlation ArBl | Fig S10B | Kolmogorov–Smirnov test | p=8.4e-26 |  |
| Population Vector Correlation AlBr | Fig S10C | Kolmogorov–Smirnov test | p=6.7e-19 |  |
| Population Vector Correlation AlBl | Fig S10D | Kolmogorov–Smirnov test | p=4.4e-15 |  |
| Spatial Correlation AB | Fig S11A | Kolmogorov–Smirnov test | p=0.0050 |  |
| Spatial Correlation AA | Fig S11B | Kolmogorov–Smirnov test | p=0.87 |  |
| Spatial Correlation ArBr | Fig S11C | Kolmogorov–Smirnov test | p=0.0054 |  |
| Spatial Correlation ArBl | Fig S11D | Kolmogorov–Smirnov test | p=0.044 |  |
| Spatial Correlation AlBr | Fig S11E | Kolmogorov–Smirnov test | p=0.047 |  |
| Spatial Correlation AlBl | Fig S11F | Kolmogorov–Smirnov test | p=0.0054 |  |

**Table S4.** Statistics used for properties of place cells from open field and linear track. Summary of statistics applied on analysis from place cells recorded during open field task (WT place cells=52 and APP-KI place cells=35) and Linear Track (WT place cells=39 and APP-KI place cells=26)

Place cells from open field and linear track
| Open field: WT Place Cells (n=52) vs APP-KI Place Cells (n=35) |  |  |  |  |
| --- | --- | --- | --- | --- |
| Linear track: WT Place Cells (n=39) vs APP-KI Place Cells (n=26) |  |  |  |  |
| Experimental measure | Figure | Test Used | P-Value | degree of freedom & F/T/z/R/ETC value |
| Open field: Field Area | Fig S8B | Wilcoxon rank sum | p=0.54 | Z=0.61 |
| Open field: # of Fields | Fig S8C | Wilcoxon rank sum | p=0.46 | Z=0.75 |
| Open field: Average Firing Rate | Fig S8D | Wilcoxon rank sum | p=0.79 | Z=0.27 |
| Open field: Peak Firing Rate | Fig S8E | Wilcoxon rank sum | p=0.94 | Z=0.074 |
| Open field: Spatial Information Index | Fig S8F | Wilcoxon rank sum | p=0.19 | Z=1.3 |
| Linear track: Field Area | Fig S8G | Wilcoxon rank sum | p=0.016 | Z=2.4 |
| Linear track: # of Fields | Fig S8H | Wilcoxon rank sum | p=0.014 | Z=2.5 |
| Linear track: Average Firing Rate | Fig S8I | Wilcoxon rank sum | p=0.88 | Z=0.15 |
| Linear track: Peak Firing Rate | Fig S8J | Wilcoxon rank sum | p=0.44 | Z=0.77 |
| Linear track: Population Vector Correlation AB | Fig S8K | Kolmogorov–Smirnov test | p=0.0099 |  |
| Linear track: Population Vector Correlation AA | Fig S8L | Kolmogorov–Smirnov test | p=0.0070 |  |
| Linear track: Spatial Correlation AB | Fig S8M | Kolmogorov–Smirnov test | p=8.7e-34 |  |
| Linear track: Spatial Correlation AA | Fig S8N | Kolmogorov–Smirnov test | p=1.3e-7 |  |

**Table S5.**
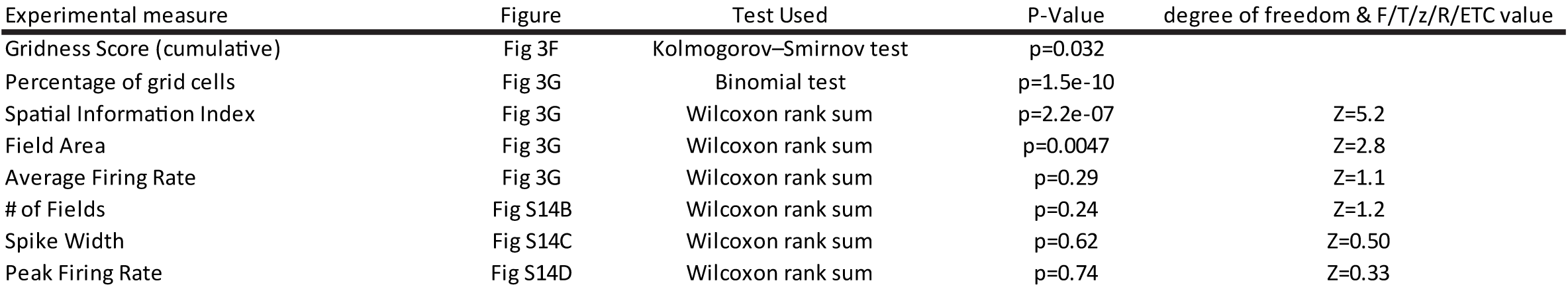
Statistics used for properties of MEC neurons from open field. Summary of statistics applied on analysis from MEC neurons recorded during open field task. (WT MEC neurons=61 and APP-KI MEC neurons=65)

**Table S6.** Statistics used for properties of MEC neurons from linear track. Summary of statistics applied on analysis from MEC neurons recorded during open field task. (MEC CA1 neurons=62 and APP-KI MEC neurons=69)

| Experimental measure | Figure | Test Used | P-Value | degree of freedom & F/T/z/R/ETC value |
| --- | --- | --- | --- | --- |
| Population Vector Correlation AB | Fig 3J | Wilcoxon rank sum | p=5.5e-27 | Z=11 |
| Population Vector Correlation AA | Fig 3J | Wilcoxon rank sum | p=0.41 | Z=0.83 |
| Field Area | Fig S16B | Wilcoxon rank sum | p=1.7e-5 | Z=4.3 |
| # of Fields | Fig S16C | Wilcoxon rank sum | p=2.9e-5 | Z=4.2 |
| Average Firing Rate | Fig S16D | Wilcoxon rank sum | p=0.48 | Z=0.70 |
| Peak Firing Rate | Fig S16E | Wilcoxon rank sum | p=0.47 | Z=0.72 |
| Population Vector Correlation AA | Fig S16F | Wilcoxon rank sum | p=0.41 | Z=0.83 |
| Population Vector Correlation AB | Fig S16G | Wilcoxon rank sum | p=5.5e-27 | Z=11 |
| Population Vector Correlation ArBr | Fig S17A | Kolmogorov–Smirnov test | p=3.0e-11 |  |
| Population Vector Correlation ArBl | Fig S17B | Kolmogorov–Smirnov test | p=4.3e-13 |  |
| Population Vector Correlation AlBr | Fig S17C | Kolmogorov–Smirnov test | p=4.0e-16 |  |
| Population Vector Correlation AlBl | Fig S17D | Kolmogorov–Smirnov test | p=1.2e-7 |  |
| Spatial Correlation AB | Fig S18A | Kolmogorov–Smirnov test | p=5.6e-5 |  |
| Spatial Correlation AA | Fig S18B | Kolmogorov–Smirnov test | p=0.51 |  |
| Spatial Correlation ArBr | Fig S18C | Kolmogorov–Smirnov test | p=8.5e-6 |  |
| Spatial Correlation ArBl | Fig S18D | Kolmogorov–Smirnov test | p=4.1e-4 |  |
| Spatial Correlation AlBr | Fig S18E | Kolmogorov–Smirnov test | p=1.8e-5 |  |
| Spatial Correlation AlBl | Fig S18F | Kolmogorov–Smirnov test | p=8.5e-6 |  |

**Table S7.** Statistics used for properties of spatially modulated MEC neurons from open field and linear track. Summary of statistics applied on analysis from spatially modulated MEC neurons recorded during open field task (WT place cells=34 and APP-KI place cells=13) and linear track (WT place cells=30 and APP-KI place cells=12)

| Experimental measure | Figure | Test Used | P-Value | degree of freedom & F/T/z/R/ETC value |
| --- | --- | --- | --- | --- |
| Spatial Information Index (cumulative) | Fig S15A | Kolmogorov–Smirnov test | p=2.2e-6 |  |
| Open field: Field Area | Fig S15C | Wilcoxon rank sum | p=0.23 | Z=1.2 |
| Open field: # of Fields | Fig S15D | Wilcoxon rank sum | p=0.84 | Z=0.21 |
| Open field: Average Firing Rate | Fig S15E | Wilcoxon rank sum | p=0.11 | Z=1.58 |
| Open field: Peak Firing Rate | Fig S15F | Wilcoxon rank sum | p=0.32 | Z=0.99 |
| Open field: Spatial Information Index | Fig S15G | Kolmogorov–Smirnov test | p=0.55 |  |
| Open field: Gridness Score | Fig S15H | Kolmogorov–Smirnov test | p=0.066 |  |
| Linear track: Field Area | Fig S15I | Wilcoxon rank sum | p=0.35 | Z=-0.93 |
| Linear track: # of Fields | Fig S15J | Wilcoxon rank sum | p=0.25 | Z=1.2 |
| Linear track: Average Firing Rate | Fig S15K | Wilcoxon rank sum | p=0.57 | Z=0.57 |
| Linear track: Peak Firing Rate | Fig S15L | Wilcoxon rank sum | p=0.43 | Z=0.79 |
| Linear track: Population Vector Correlation AB | Fig S15M | Kolmogorov–Smirnov test | p=1.7e-4 |  |
| Linear track: Population Vector Correlation AA | Fig S15N | Kolmogorov–Smirnov test | p=0.15 |  |
| Linear track: Spatial Correlation AB | Fig S15O | Kolmogorov–Smirnov test | p=0.43 |  |
| Linear track: Spatial Correlation AA | Fig S15P | Kolmogorov–Smirnov test | p=0.12 |  |

**Table S8.** Statistics used for LFP analysis on CA1. Summary of statistics applied on LFP analysis from CA1 recording during open field task

LFP analyses: CA1
| Experimental measure | Figure | Test Used | Sample | P-Value | degree of freedom & F/T/z/R/ETC value |
| --- | --- | --- | --- | --- | --- |
| Normalized oscillatory power (Slow and Fast gamma) | Fig 2C | unpaired t-test with FDR corrected for 100 multiple comparison | WT=10 mice, APP-KI = 10 mice | q=0.023 (average) |  |
| Occurrence rate of fast gamma | Fig 2E | Wilcoxon rank sum | WT=10 mice, APP-KI = 10 mice | P= 1.8e-4 | Z=3.74 |
| Occurrence rate of slow gamma | Fig 2E | Wilcoxon rank sum | WT=10 mice, APP-KI = 10 mice | P=0.38 | Z=0.87 |
| Theta to slow gamma coupling | Fig 2G | Wilcoxon rank sum | WT=10 mice, APP-KI = 10 mice | p=0.35 | Z=0.95 |
| Theta to fast gamma coupling | Fig 2G | Wilcoxon rank sum | WT=10 mice, APP-KI = 10 mice | p=0.045 | Z=2.00 |
| Phase locking of spikes to slow gamma | Fig 2I | Wilcoxon rank sum | WT=104 neurons, APP-KI=105 neurons | p=0.40 | Z=0.84 |
| Phase locking of spikes to fast gamma | Fig 2I | Wilcoxon rank sum | WT=104 neurons, APP-KI=105 neurons | p=8.5e-10 | Z=6.13 |
| Phase locking of spikes to fast gamma | Fig S20A | binomial test | WT=(64/104),APP-KI=(9/105) | p=1.5e-12 |  |
| Phase locking of spikes to slow gamma | Fig S20B | binomial test | WT=(48/104),APP-KI=(38/105) | p=0.041 |  |

**Table S9.**
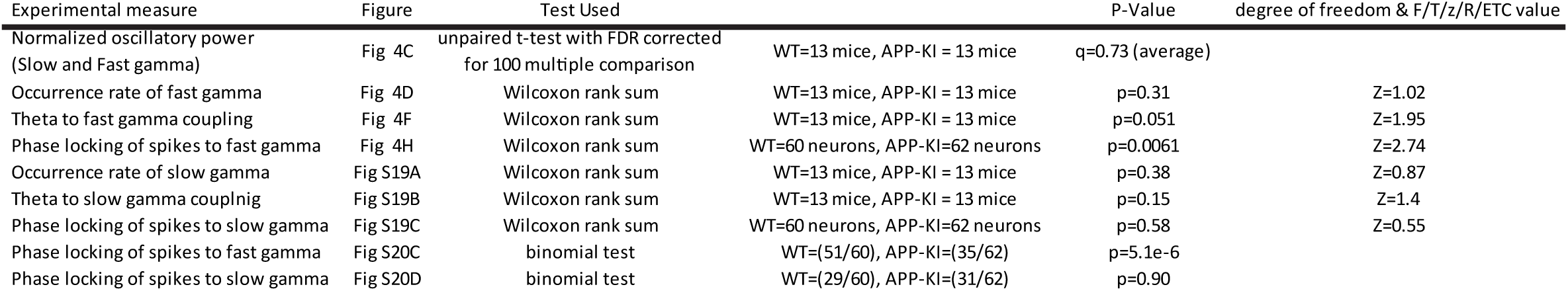
Statistics used for LFP analysis on MEC. Summary of statistics applied on LFP analysis from MEC recording during open field task

**Table S10.**
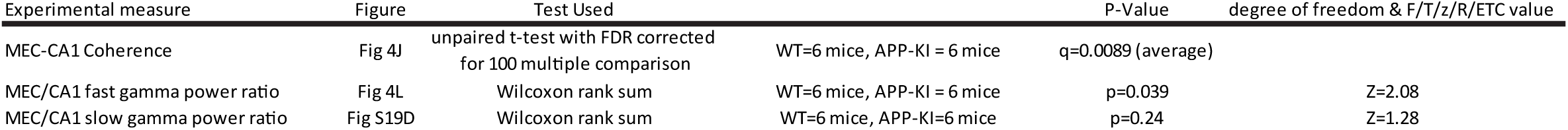
Statistics used for LFP analysis on CA1-MEC. Summary of statistics applied on LFP analysis from simultaneous recording of CA1 and MEC regions during open field task

**Table S11.** Statistics used for spatial tuning vs spatial correlation AB. Summary of statistics applied on spatial tuning vs spatial correlation AB analysis

Correlation between spatial tuning and remapping
| Experimental measure | Figure | Test Used | Sample # | P-Value | degree of freedom & F/T/z/R/ETC value |
| --- | --- | --- | --- | --- | --- |
| Spatial Tuning vs Spatial Correlation (WT CA1) | Fig 3K | Pearson Correlation | WT=105 neurons | p=0.14 | r(105)=-0.15 |
| Spatial Tuning vs Spatial Correlation (WT MEC) | Fig 3K | Pearson Correlation | WT=61 neurons | p=0.09 | r(61)=-0.22 |
| Spatial Tuning vs Spatial Correlation (APP-KI CA1) | Fig 3K | Pearson Correlation | APP-KI=113 neurons | p=0.22 | r(113)=-0.14 |
| Spatial Tuning vs Spatial Correlation (APP-KI MEC) | Fig 3K | Pearson Correlation | APP-KI=65 neurons | p=1e-04 | r(66)=-0.45 |

**Table S12.** Statistics used for spatial tuning and spatial correlation AB across age. Summary of statistics applied on spatial tuning and spatial correlation AB across age.

Spatial tuning and remapping across age
| Experimental measure | Figure | Test Used | Sample # | P-Value | degree of freedom & F/T/z/R/ETC value |
| --- | --- | --- | --- | --- | --- |
| Spatial Information across age (CA1) | Fig S4A | Pearson Correlation | WT=105, APP-KI=113 | p(WT)=0.13, p(APP-KI)=0.58 | r(105,WT)= -0.15 , r(116,APP-KI)=0.053 |
| Spatial Information across age (MEC) | Fig S4B | Pearson Correlation | WT=61,APP-KI=65 | p(WT)=0.25, p(APP-KI)=0.26 | r(61,WT)= 0.15 , r(66,APP-KI)=0.14 |
| Spatial Correlation AB across age (CA1) | Fig S4C | Pearson Correlation | WT=105, APP-KI=113 | p(WT)=0.21, p(APP-KI)=0.21 | r(105,WT)= 0.13 , r(113,APP-KI)=0.14 |
| Spatial Correlation AB across age (MEC) | Fig S4D | Pearson Correlation | WT=61,APP-KI=65 | p(WT)=0.017, p(APP-KI)=0.39 | r(61,WT)= -0.30 , r(65,APP-KI)=0.11 |

**Table S13.**
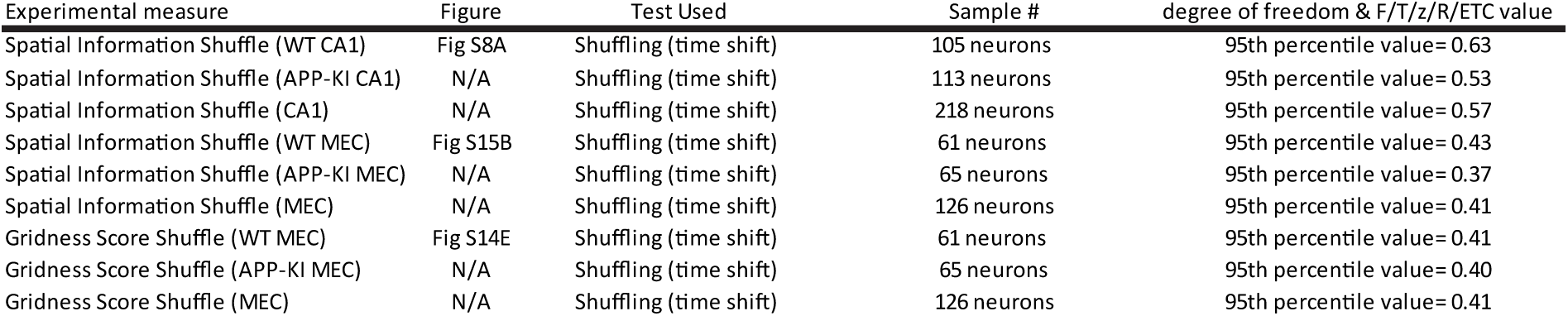
Shuffled values. Summary of 95^th^ percentile shuffled values obtained for spatial information and gridness score.

Shuffled values
| Experimental measure | Figure | Test Used | Sample # | degree of freedom & F/T/z/R/etc value |
| --- | --- | --- | --- | --- |
| Spatial Information Shuffle (WT CA1) | Fig S8A | Shuffling (time shift) | 105 neurons | 95th percentile value= 0.63 |
| Spatial Information Shuffle (APP-KI CA1) | N/A | Shuffling (time shift) | 113 neurons | 95th percentile value= 0.53 |
| Spatial Information Shuffle (CA1) | N/A | Shuffling (time shift) | 218 neurons | 95th percentile value= 0.57 |
| Spatial Information Shuffle (WT MEC) | Fig S15B | Shuffling (time shift) | 61 neurons | 95th percentile value= 0.43 |
| Spatial Information Shuffle (APP-KI MEC) | N/A | Shuffling (time shift) | 65 neurons | 95th percentile value= 0.37 |
| Spatial Information Shuffle (MEC) | N/A | Shuffling (time shift) | 126 neurons | 95th percentile value= 0.41 |
| Gridness Score Shuffle (WT MEC) | Fig S14E | Shuffling (time shift) | 61 neurons | 95th percentile value= 0.41 |
| Gridness Score Shuffle (APP-KI MEC) | N/A | Shuffling (time shift) | 65 neurons | 95th percentile value= 0.40 |
| Gridness Score Shuffle (MEC) | N/A | Shuffling (time shift) | 126 neurons | 95th percentile value= 0.41 |

**Table S14.**
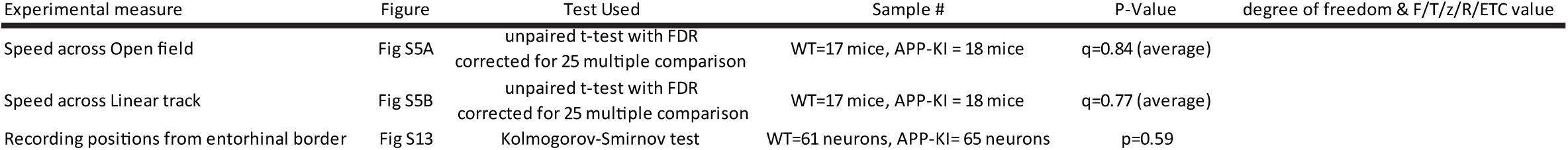
Statistics used for animal Behavior and Recording Positions. Summary of statistics applied on speed analysis and recording positions from entorhinal border.

**Table S15.** Statistics used for Principal neuron classification. Summary of statistics applied on peak-valley duration obtained from CA1 neurons (WT=138 neurons and APP-KI=164 neurons) and MEC neurons (WT=81 neurons and APP-KI=99 neurons)

Principal neuron classification
| Experimental measure | Figure | Test Used | Sample # | P-Value | degree of freedom & F/T/z/R/etc value |
| --- | --- | --- | --- | --- | --- |
| CA1 neurons Peak-Valley Duration | Fig S6A | Kolmogorov-Smirnov test | WT=138 neurons, APP-KI= 164 neurons | 0.17 |  |
| MEC neurons Peak-Valley Duration | Fig S6B | Kolmogorov-Smirnov test | WT=81 neurons, APP-KI= 99 neurons | 0.33 |  |

